# The Tryptophan Metabolizing Enzyme Indoleamine 2,3-Dioxygenase 1 Regulates Polycystic Kidney Disease Progression

**DOI:** 10.1101/2022.05.04.490641

**Authors:** Dustin T. Nguyen, Emily K. Kleczko, Nidhi Dwivedi, Berenice Y. Gitomer, Michel B. Chonchol, Eric T. Clambey, Raphael A. Nemenoff, Jelena Klawitter, Katharina Hopp

## Abstract

Autosomal dominant polycystic kidney disease (ADPKD), the most common monogenic nephropathy, is characterized by phenotypic variability exceeding genic effects. Dysregulated metabolism and immune cell function are key disease modulators. The tryptophan metabolites, kynurenines, produced through IDO1, are known immunomodulators. Here, we study the role of tryptophan metabolism in PKD using an orthologous disease model (C57Bl/6J *Pkd1*^RC/RC^). We found elevated kynurenine and IDO1 levels in *Pkd1*^RC/RC^ kidneys versus wildtype. Further, IDO1 levels were increased in ADPKD cell lines and patient cyst cells. Genetic *Ido1* loss in *Pkd1*^RC/RC^ animals resulted in reduced PKD severity as measured by %kidney weight/body weight and cystic index. Consistent with a immunomodulatory role of kynurenines, *Pkd1*^RC/RC^;*Ido1*^-/-^ mice presented with significant changes in the cystic immune microenvironment (CME) versus controls. Of note, kidney macrophage numbers decreased and CD8^+^ T cell numbers increased, both known PKD modulators. Also, pharmacological IDO1 inhibition using a tryptophan analog in *Pkd1*^RC/RC^ animals resulted in less severe PKD versus controls with similar changes in the CME as in the genetic model. Together, our data suggest that tryptophan metabolism is dysregulated in ADPKD and that its inhibition results in changes to the CME and slows disease progression, making IDO1 a novel therapeutic target for ADPKD.

## Introduction

Autosomal dominant polycystic kidney disease (ADPKD) is the most common, life threatening genetic kidney disease(1, 2). It is characterized by progressive kidney cyst growth leading to organ failure and accounts for 5-10% of end stage kidney disease cases worldwide(3, 4). Tolvaptan, a vasopressin receptor antagonist, is the only FDA approved therapy for ADPKD, which slows cyst growth but also impairs quality of life, underscoring the need for alternative therapies(5, 6). ADPKD is predominantly caused by mutations to *PKD1* or *PKD2;* but manifests with significant inter- and intrafamilial phenotypic variability that exceeds genic or allelic effects(7–9). Disease variability has been linked to genetic modifiers, kidney injury, and environmental/lifestyle factors(7, 10–12).

Accumulating evidence suggest that metabolic reprogramming is a key feature of ADPKD(13–15). In the HALT-PKD Study A population, overweight/obesity were shown to be strong independent risk factors of total kidney volume increase and eGFR decline(10). Correlatively, preclinical studies in PKD models have shown that mild-to-moderate caloric restriction, time restricted feeding, or ketogenic diet ameliorate kidney cyst growth(16–18). Indeed, targeting specific metabolic pathways found to be dysregulated in PKD such as glycolysis, fatty acid oxidation, or arginine, glutamine, and methionine metabolism alleviated cystic kidney disease in PKD models(19–24).

Data from our group and others suggest that tryptophan metabolism may also play a role in ADPKD progression(25, 26). Tryptophan is catabolized to kynurenine via tryptophan 2,3-dioxygenase (TDO) or indoleamine 2,3-dioxygnease (IDO1, IDO2, **Supplemental Figure 1**). While TDO is predominantly expressed in the liver, IDO1/2 are expressed in kidney epithelial- and immune cells, with IDO1 having higher catalytic activity for tryptophan(27–29). Kynurenine and/or its metabolite, kynurenic acid, are known drivers of oxidative stress, dysregulated calcium homeostasis, and mitochondrial dysfunction(30–32). As such, they are considered uremic toxins; their serum levels correlate with chronic kidney disease (CKD) severity(33). Interestingly, serum metabolomics of participants of the Modification of Diet in Renal Disease Study showed that levels of kynurenic acid were selectively elevated in ADPKD patients compared to other CKD patients. Also, we recently published that ADPKD patients have higher plasma kynurenine concentrations compared to healthy subjects and levels further increased with disease progression(25, 26).

In cancer, a disease with multiple parallels to PKD, plasma kynurenine levels and tumor expression of IDO1 correlate with cancer survival and clinical outcome(34–38). Elevated kynurenine levels and IDO1 upregulation promote an immunosuppressive microenvironment and thus allow tumor escape from immune destruction. This includes suppression of anti-tumorigenic CD8^+^ T cells through upregulation of the immune checkpoint PD-1/PD-L1, generation of pro-tumorigenic regulatory T cells (TReg), and promotion of tumor-associated M2 macrophage polarization(39, 40). Indeed, inhibition of tryptophan metabolism via IDO1 inhibitors has been FDA approved for multiple cancers either as mono- or combination therapy with anti-PD-1(38). Interestingly, recent data by others and us suggest that both innate and adaptive immune cells, such as M2-like macrophages and CD8^+^ T cells, are also important modulators of kidney cyst growth in PKD murine models(41–43). However, the functional impact of the tryptophan pathway on cyst growth and its potential as a therapeutic target in ADPKD has not been clearly established.

Here, we utilize an orthologous ADPKD1 model to study tryptophan metabolism in PKD(43–45). We found both tryptophan metabolites and IDO1 expression to be elevated in ADPKD1 mice correlative with disease progression. Genetic loss and pharmaceutical inhibition of IDO1 slowed cyst growth in our model. This was associated with reduced numbers of kidney M2-like resident macrophages and TRegs, reduced expression of PD-1/PD-L1, and an increase of CD8^+^ T cell numbers within the adaptive immune cell population. Together, our data implicates tryptophan metabolism as a novel modifier of ADPKD progression and suggests that FDA approved IDO1 inhibitors may represent a new treatment approach for ADPKD.

## Results

### Tryptophan metabolism via IDO1 is dysregulated in Pkd1^RC/RC^ mice

To determine whether tryptophan metabolism is abnormally regulated in murine PKD, we performed metabolomic analyses of metabolites within the tryptophan pathway using the orthologous ADPKD model C57Bl/6J *Pkd1* p.R3277C (*Pkd1*^RC/RC^)(43–45). Comparing kidney metabolites of strain, age, and gender matched wildtype (WT) mice, we found that tryptophan levels remained stable throughout PKD progression (3 to 9 months [mo]) and comparable to WT levels (**Table 1, Supplemental Figure 1**). However, the immunosuppressive metabolites kynurenine/kynurenic acid were significantly upregulated in *Pkd1*^RC/RC^ kidneys correlative with disease severity(46). Similarly, levels of xanthurenic acid, a kynurenic acid metabolite, were elevated compared to WT. Consistent with increased production of kynurenic and xanthurenic acid, *Pkd1*^RC/RC^ kidneys displayed significantly decreased levels of picolinic acid compared to WT (**Table 1**). Picolinic acid is an isomer of nicotinic acid, a derivative of nicotinamide, which was also significantly decreased and has been shown to slow cyst growth and improve kidney function in two PKD models(47). No changes were observed in levels of anthranilic acid and quinolinic acid (data not shown). Overall, these data highlight that tryptophan metabolism is dysregulated in *Pkd1*^RC/RC^ kidneys with a shift towards the production of known immunosuppressive metabolites.

**Table 1.** Levels of tryptophan metabolites in *Pkd1*^RC/RC^ kidneys.

| Metabolite (Abundance/mg tissue [normalized]) |  |  |  |  |  |  |
| --- | --- | --- | --- | --- | --- | --- |
|  | Tryptophan | Kynurenine | Kynurenic acid | Xanthurenic acid | Picolinic acid | Nicotinamide |
| **3 months** |  |  |  |  |  |  |
| WT | 2.23x10<sup>-2</sup>±0.816x10<sup>-2</sup> | 1.96x10<sup>-4</sup>±0.804x10<sup>-4</sup> | 1.10x10<sup>-2</sup>±0.476x10<sup>-2</sup> | 2.92x10<sup>-2</sup>±0.766x10<sup>-2</sup> | 6.78x10<sup>-3</sup>±2.11x10<sup>-3</sup> | 3.86±0.995 |
| *Pkd1<sup>RC/RC</sup>* | 1.76x10<sup>-2</sup>±1.02x10<sup>-2</sup> | 10.6x10<sup>-4</sup>±9.49x10<sup>-4</sup> | 6.34x10<sup>-2</sup>±3.97x10<sup>-2**</sup> | 13.0x10<sup>-2</sup>±8.52x10<sup>-2</sup> | 4.70x10<sup>-3</sup>±2.54x10<sup>-3</sup> | 2.78±1.25* |
| Fold change | 0.789 | 5.41 | 5.76 | 4.45 | 0.693 | 0.720 |
| **6 months** |  |  |  |  |  |  |
| WT | 2.36x10<sup>-2</sup>±0.759x10<sup>-2</sup> | 2.95x10<sup>-4</sup>±1.69x10<sup>-4</sup> | 0.735x10<sup>-2</sup>±0.291x10<sup>-2</sup> | 4.17x10<sup>-2</sup>±3.42x10<sup>-2</sup> | 6.07x10<sup>-3</sup>±1.20x10<sup>-3</sup> | 3.46±1.02 |
| *Pkd1<sup>RC/RC</sup>* | 2.02x10<sup>-2</sup>±0.750x10<sup>-2</sup> | 14.0x10<sup>-4</sup>±7.29x10<sup>-4*</sup> | 9.70x10<sup>-2</sup>±4.39x10<sup>-2****</sup> | 19.6x10<sup>-2</sup>±10.5x10<sup>-2***</sup> | 3.23x10<sup>-3</sup>±0.672x10<sup>-3*</sup> | 2.07±0.470** |
| Fold change | 0.856 | 4.75 | 13.2 | 4.70 | 0.532 | 0.598 |
| **9 months** |  |  |  |  |  |  |
| WT | 1.94x10<sup>-2</sup>±0.629x10<sup>-2</sup> | 2.87x10<sup>-4</sup>±2.43x10<sup>-4</sup> | 0.688x10<sup>-2</sup>±0.371x10<sup>-2</sup> | 1.59x10<sup>-2</sup>±0.600x10<sup>-2</sup> | 7.58x10<sup>-3</sup>±2.17x10<sup>-3</sup> | 3.61±1.22 |
| *Pkd1<sup>RC/RC</sup>* | 1.73x10<sup>-2</sup>±0.637x10<sup>-2</sup> | 21.3x10<sup>-4</sup>±12.2x10<sup>-4**** #</sup> | 8.53x10<sup>-2</sup>±3.15x10<sup>-2****</sup> | 22.3x10<sup>-2</sup>±12.3x10<sup>-2****</sup> | 4.13x10<sup>-3</sup>±2.03x10<sup>-3**</sup> | 2.54±1.19 |
| Fold change | 0.891 | 7.42 | 12.4 | 14.0 | 0.545 | 0.704 |

N = 5 males, 5 females per genotype and time point; Fold change: *Pkd1<sup>RC/RC</sup>* versus WT; P-values: *Pkd1<sup>RC/RC</sup>* versus wildtype \*<0.05, \*\*<0.01, \*\*\*<0.001, \*\*\*\*<0.0001; 3 months *Pkd1<sup>RC/RC</sup>* versus 9 months *Pkd1<sup>RC/RC</sup>* #<0.05

Western blot analysis of whole kidney homogenates showed increased expression of IDO1 in kidneys of *Pkd1*^RC/RC^ mice compared to WT (**Figure 1A**). Similar increases were observed in 9-12 cells *(PKD1*^-/-^) compared to renal cortical epithelial cells (RCTE; *PKD1*^+/+^, **Figure 1B**). The expression levels of IDO1 could further be amplified by stimulating RCTE or 9-12 cells with interferon gamma (IFNγ), as has been reported in the literature and which we have shown to be elevated in *Pkd1*^RC/RC^ kidneys (**Figure 1B**)(43, 48). Further, we found high levels of IDO1 in cyst lining epithelial kidney cells derived from ADPKD patients (**Figure 1C**). We confirmed elevated levels of IDO1 by immunofluorescence (**Figure 1D**). In WT mice, IDO1 expression was sparsely detected within interstitial cells and not in the kidney epithelium. However, *Pkd1*^RC/RC^ mice showed increased levels of IDO1 expression in cyst lining- and interstitial cells. Based on the literature, the interstitial cells expressing IDO1 are likely macrophages or dendritic cells, which have been shown to upregulate IDO1 in various types of cancer resulting in increased tumorigenesis(36, 49). These data suggest that the observed dysregulation of tryptophan catabolism may be attributed to overexpression of IDO1 within the cystic epithelia as well as in kidney immune cells.

**Figure 1.**
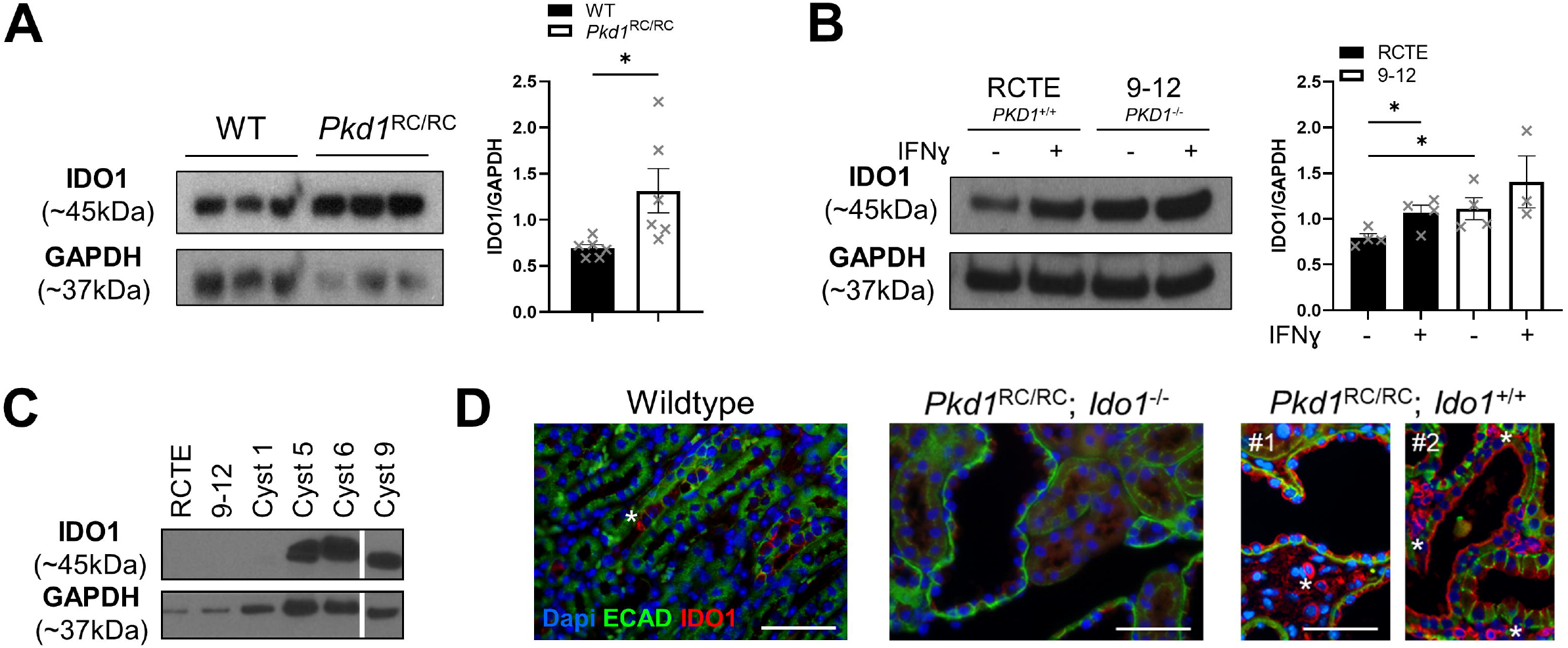
*Pkd1*^RC/RC^ animals present with overexpression of IDO1. (A) Western blot probing for IDO1 (left) and quantification (right) using wildtype (WT) and *Pkd1*^RC/RC^ kidney homogenates, highlighting upregulation of IDO1 in *Pkd1*^RC/RC^ kidney compared to WT. (Quantification: N= 3males/3females, 9mo). (B) Western blot probing for IDO1 (left) and quantification (right) of cell lysates obtained from normal renal cortical epithelial cells (RCTE, WT for *PKD1*) or 9-12 cells (null for *PKD1)* ± IFNγ stimulation, confirming overexpression of IDO1 in PKD-relevant human cell lines compared to control. IDO1 expression levels were further increased by the cytokine IFNγ, which is known to be upregulated in PKD kidneys (N=4 per condition). (C) Western blot probing for IDO1 levels in epithelial cells obtained from individual cysts of ADPKD patients (each cyst was derived from a different patient, PKD genotype unknown). Most tested cysts show very high levels of IDO1; relative to IDO1 levels in RCTE or 9-12 cells (same level of protein was loaded, but insufficient exposure time to detect IDO1 in RCTE or 9-12 cells). This provides direct clinical relevance for dysregulation of the tryptophan pathway in ADPKD patient’ kidneys. (D) Immunofluorescence images probing for IDO1 (red), e-cadherin (ECAD, green, epithelial cells), and DAPI (blue, nuclei). IDO1 is sparsely expressed in wildtype kidneys but upregulated in kidney cystic epithelial cells and interstitial cells of *Pkd1*^RC/RC^ kidneys. *Ido1* knock-out animals served as negative control. *references IDO1 positive interstitial cells in WT or *Pkd1*^RC/RC^; *Ido1*^+/+^ kidneys, #1 and #2 references two different *Pkd1*^RC/RC^; *Ido1*^+/+^ animals. Scale bar = 50μm. Statistics: Graphs, mean ± SEM; Analyses, unpaired t test (A, B). P *<0.05, comparisons with non-significant statistics are not shown.

### Genetic loss of Ido1 alleviates PKD severity and corrects tryptophan metabolism abnormalities

To functionally establish a role for tryptophan metabolism and IDO1 in PKD pathogenesis, we crossed C57Bl/6J *Pkd1*^RC/RC^ mice with C57Bl/6J *Ido1*^-/-^ mice. We aged the resulting second filial generation animals (*Pkd1*^RC/RC^;*Ido1*^+/+^ or *Pkd1^RC/RC^;Ido1^-/-^*) to 3mo or 6mo of age and evaluated histopathologicalIphysiological features commonly analyzed in murine PKD studies (**Figure 2, Supplemental Figure 2**). At 3mo of age we observed no difference in gross histological appearance, percent kidney weight normalized to body weight (%KW/BW), or cystic volume/index between *Pkd1*^RC/RC^;*Ido1*^+/+^ or *Pkd1^RC/RC^;Ido1^-/-^* mice (**Figure 2A-C, Supplemental Figure 2A**). However, at 6mo of age *Pkd1^RC/RC^;Ido1^-/-^* mice presented with decreased cystic kidney disease compared to *Pkd1*^RC/RC^;*Ido1*^+/+^ mice (**Figure 2A-C, Supplemental Figure 2A**). Indeed, it appears that cyst growth from 3 to 6mo of age in *Pkd1*^RC/RC^;*Ido1*^-/-^ mice was halted compared to controls, as evidenced by a significant “reduction” in kidney cyst count and size (**Figure 2C**). We did not detect a difference in fibrotic burden or kidney function (BUN levels, **Supplemental Figure 2B, C**).

**Figure 2.**
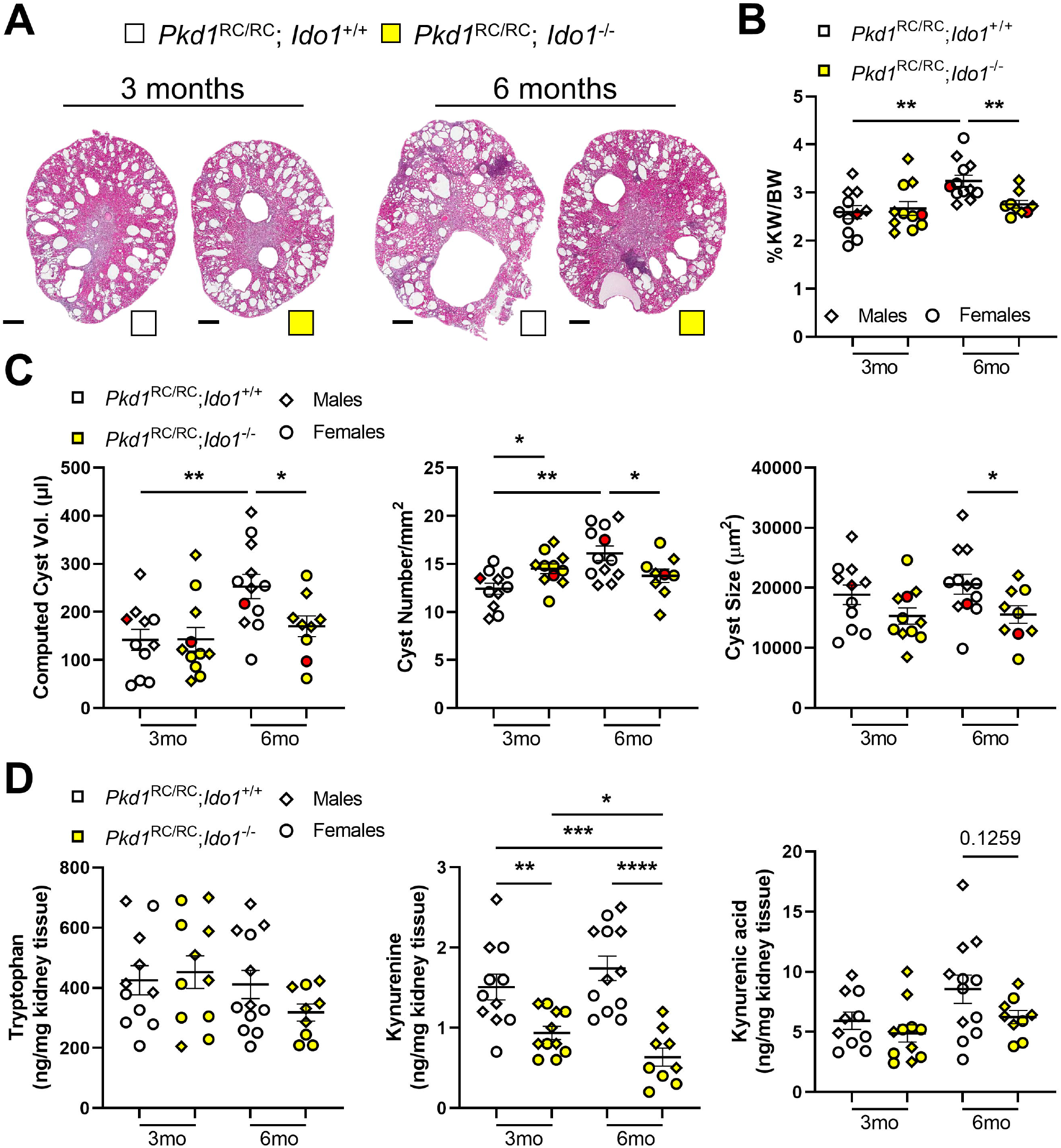
Genetic loss of *Ido1* slows cyst growth and reduces tryptophan catabolite levels in kidneys of an orthologous ADPKD1 model. (A) Kidney H&E cross sections of 3mo and 6mo old *Pkd1*^RC/RC^; *Ido1^+/+^* (white) and *Pkd1*^RC/RC^; *Ido1*^-/-0^ (yellow) kidneys showing overall decreased cystic disease severity in 6mo old PKD *Ido1* null versus *Ido1* wildtype (WT) animals. Scale bar: 500μm. Quantification of (B) %KW/BW, (C) computed cyst volume (cystic index [Supplemental Figure 2A] multiplied by kidney weight), cyst number, and cyst size in *Pkd1*^RC/RC^; *Ido1*^+/+^ (white) and *Pkd1*^RC/RC^; *Ido1*^-/-^ (yellow) animals, together providing statistical significance for reduced PKD severity in 6mo old PKD *Ido1* null versus *Ido1* WT animals. Red data point depicts the animal shown in (A). (D) Quantification of significantly altered tryptophan catabolites assayed via mass spectrometry in *Pkd1*^RC/RC^; *Ido1^+/+^* (white) and *Pkd1*^RC/RC^; *Ido1*^-/-^ (yellow) kidneys. Loss of *Ido1* partially corrected the observed increased levels of the immunosuppressive tryptophan catabolites, kynurenine and kynurenic acid, seen in *Pkd1*^RC/RC^ kidneys (Supplemental Figure 2D). N= 5males (diamond), 4-7females (circle) per genotype and time point. Statistics: Graphs: mean ± SEM; Analyses: one-way ANOVA with Tukey’s multiple comparison test. P *<0.05, **<0.01, ***<0.001, ****<0.0001, comparisons with non-significant statistics are not shown.

Consistent with *Ido1* loss, we observed significantly decreased levels of the immunosuppressive metabolite kynurenine in *Pkd1*^RC/RC^;*Ido1*^-/-^ animals compared to controls (**Figure 2D**). Indeed, the kynurenine levels of 6mo old *Pkd1*^RC/RC^;*Ido1*^-/-^ mice were comparable to WT levels (**Supplemental Figure 2D**). Neither tryptophan nor kynurenic acid were significantly altered between the two *Ido1* genotypes in the setting of PKD, but we observed a clear trend towards decreased kynurenic acid levels at 6mo of age in *Pkd1*^RC/RC^;*Ido1*^-/-^ animals versus controls (**Figure 2D**).

### Ido1 loss is associated with a CME favorable to halt cyst progression

We compared kidney immune cell types of *Pkd1*^RC/RC^;*Ido1*^+/+^ and *Pkd1*^RC/RC^;*Ido1*^-/-^ mice at 3mo and 6mo of age using flow cytometry of kidney single cell suspensions (**Figure 3, Supplemental Figure 3**)(41, 43, 50). Overall, we found that the observed decreased PKD severity in *Pkd1*^RC/RC^;*Ido1*^-/-^ mice compared to *Pkd1*^RC/RC^*Ido1*^+/+^ mice at 6mo of age was associated with lower numbers of kidney immune cells (CD45^+^, **Figure 3A**) and a significant decrease in infiltrating (F4/80^lo^; CD11b^+^) and resident (F4/80^hi^; CD11b^+^) macrophages, neutrophils (GR1^+^), dendritic cells (DCs, CD11b^+^; CD11c^+^) and natural killer cells (NK cells, NKp46^+^) compared to control (**Figure 3B, Supplemental Figure 3A-C**). While less is known about the role of neutrophils, DCs, or NK cells in PKD, ample evidence supports a role of “M2-like” macrophages driving kidney/liver cysts growth in murine PKD models(41, 42, 51–57). With respect to adaptive immune cells, we did not find a change in overall T cell (TCRβ^+^, CD4^+^, or CD8^+^) numbers associated with the less serve disease observed in *Pkd1*^RC/RC^;*Ido1*^-/-^ mice compared to *Pkd1*^RC/RC^;*Ido1*^+/+^ mice at 6mo of age (**Figure 3C**). However, the distribution of T cell subtypes changed, with 6mo old *Pkd1*^RC/RC^*Ido1*^-/-^ mice having significantly more CD8^+^ T cells compared to control (**Figure 3D**). Given our previously published findings that CD8^+^ T cells play an anti-cystogenic role in the *Pkd1*^RC/RC^ model, this increase in CD8^+^ T cell numbers supports a reduction in cyst severity in PKD *Ido1*^-/-^ animals compared to control(43). This increase of CD8^+^ T cells is counterbalanced by a decrease in double negative T cells, whose functional role has not been studied in PKD, but we and others have reported that their numbers increase in ADPKD compared to control in *Pkd1*^RC/RC^ kidneys as well as ADPKD patient kidneys (**Supplemental Figure 3D**)(50, 58).

**Figure 3.**
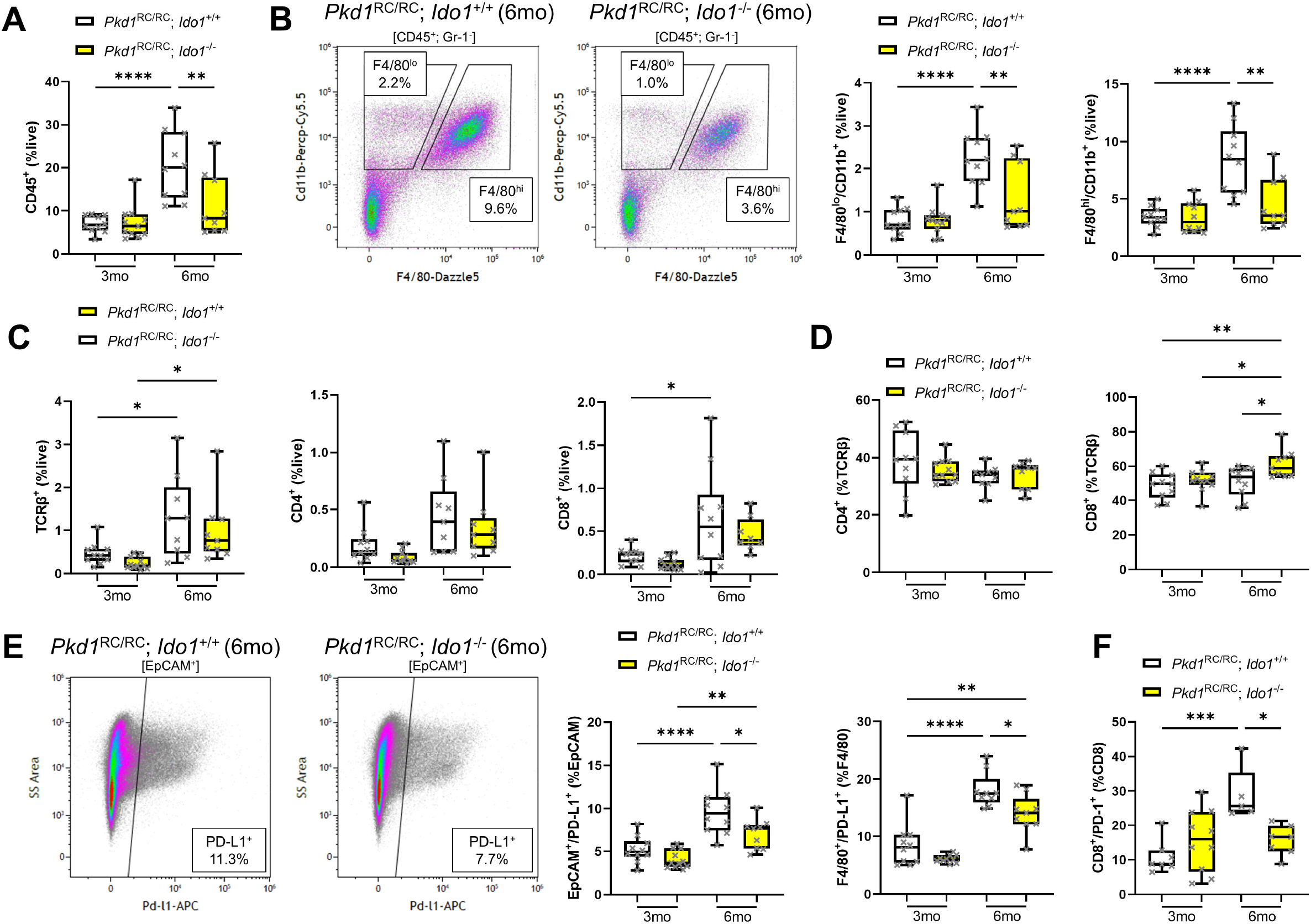
The kidney immune landscape of *Pkd1*^RC/RC^; Ido1^-/-^ mice favors slowed PKD progression. *Pkd1*^RC/RC^; *Ido1*^+/+^ (white), *Pkd1*^RC/RC^; *Ido1*^-/-^ (yellow); (A) Flow cytometry quantification of CD45^+^ immune cells within single cell suspensions of *Pkd1*^RC/RC^; *Ido1^+/+^* and *Pkd1*^RC/RC^; *Ido1*^-/-^ kidneys showing an increase of CD45^+^ cells with PKD progression (3-6mo of age, *Pkd1*^RC/RC^; *Ido1*^+/+^ mice) and a decrease in PKD *Ido1* null versus wildtype (WT) animals at 6mo of age when reduced PKD severity was observed. (B) Representative flow cytometry plot indicating the gating strategy of infiltrating (F4180^lo^; CD11b^+^) versus resident (F4/80^hi^; CD11b^+^) macrophages (left), and quantification (right), highlighting a significant increase of macrophages as disease progresses from 3-6mo of age in *Pkd1*^RC/RC^; *Ido1*^+/+^ mice and a decrease at 6mo of age in *Pkd1*^RC/RC^; *Ido1*^-/-^ mice versus WT. (C) Flow cytometry data quantification of all T cells (TCRβ^+^) and CD4^+^ or CD8^+^ subpopulations. The numbers of CD4^+^ or CD8^+^ cells in kidneys did not change significantly upon *Ido1* loss independent of disease severity. (D) Quantification of CD4^+^ and CD8^+^ T cell numbers as %TCRβ^+^ cells, showing a shift in distribution of T cell subpopulations with an increase in CD8^+^ T cell numbers upon *Ido1* loss and reduced PKD severity (6mo of age). (E) Representative flow cytometry plot (left) and quantification (right) of immune checkpoint ligand PD-L1 expression on kidney epithelial cells (plot and quantification, EpCAM^+^) and macrophages (quantification only, F4/80^+^) indicating reduced expression in 6mo old *Pkd1*^RC/RC^; *Ido1*^-/-^ animals versus control (*Pkd1*^RC/RC^; *Ido1*^+/+^). (F) Quantification of immune checkpoint receptor PD-1 expression on CD8^+^ T cells showing a decrease in expression in 6mo old *Pkd1*^RC/RC^; *Ido1*^-/-^ animals versus control (*Pkd1*^RC/RC^; *Ido1*^+/+^). Statistics: Graphs: box plot, whiskers 10-90^th^ percentile; Analyses: one-way ANOVA with Tukey’s multiple comparison test. P *<0.05, **<0.01, ***<0.001, ****<0.0001, comparisons with non-significant statistics are not shown. N= 5males, 4-7females per genotype/time point.

Kynurenines have been reported in the cancer literature to support immune escape of tumors via engagement of the PD-1/PD-L1 immune checkpoint and differentiation of CD4^+^ T cells into immunosuppressive TRegs(59, 60). In PKD, we found activation of the PD-1/PD-L1 checkpoint as well as increased numbers of T_Regs_ (CD4^+^/FoxP3^+^) in *Pkd1*^RC/RC^ kidneys correlative to disease progression (**Supplemental Figure 4**, unpublished)(61). We observed a downward trend in kidney TReg numbers in *Pkd1*^RC/RC^;*Ido1*^-/-^ animals compared to control and a significant decrease in PD-L1 expression on the kidney epithelium (EpCAM^+^/PD-L1^+^), macrophages (F4/80^+^/PD-L1^+^), and PD-1 on CD8^+^ T cells at 6mo of age (**Figure 3E, F, Supplemental Figure 3E**).

### Pharmaceutical inhibition of IDO1 slows cyst growth in Pkd1^RC/RC^ mice and changes the CME towards an anti-cystogenic composition

We treated 1mo C57Bl/6J *Pkd1*^RC/RC^ mice with 400mg/kg 1-Methyl-D-tryptophan (1-MT) via gavage twice daily for 3 weeks. 1 -MT is a synthetic tryptophan analog acting as a competitive inhibitor(62, 63). *Pkd1*^RC/RC^ mice treated with 1-MT versus control displayed significantly less severe PKD as apparent by histology, %KW/BW, cyst volume/index, and cyst number (**Figure 4A-C, Supplemental Figure 5A**). Treatment did not impact cyst size nor fibrotic burden or kidney function (**Figure 4A-C, Supplemental Figure 5A**).

**Figure 4.**
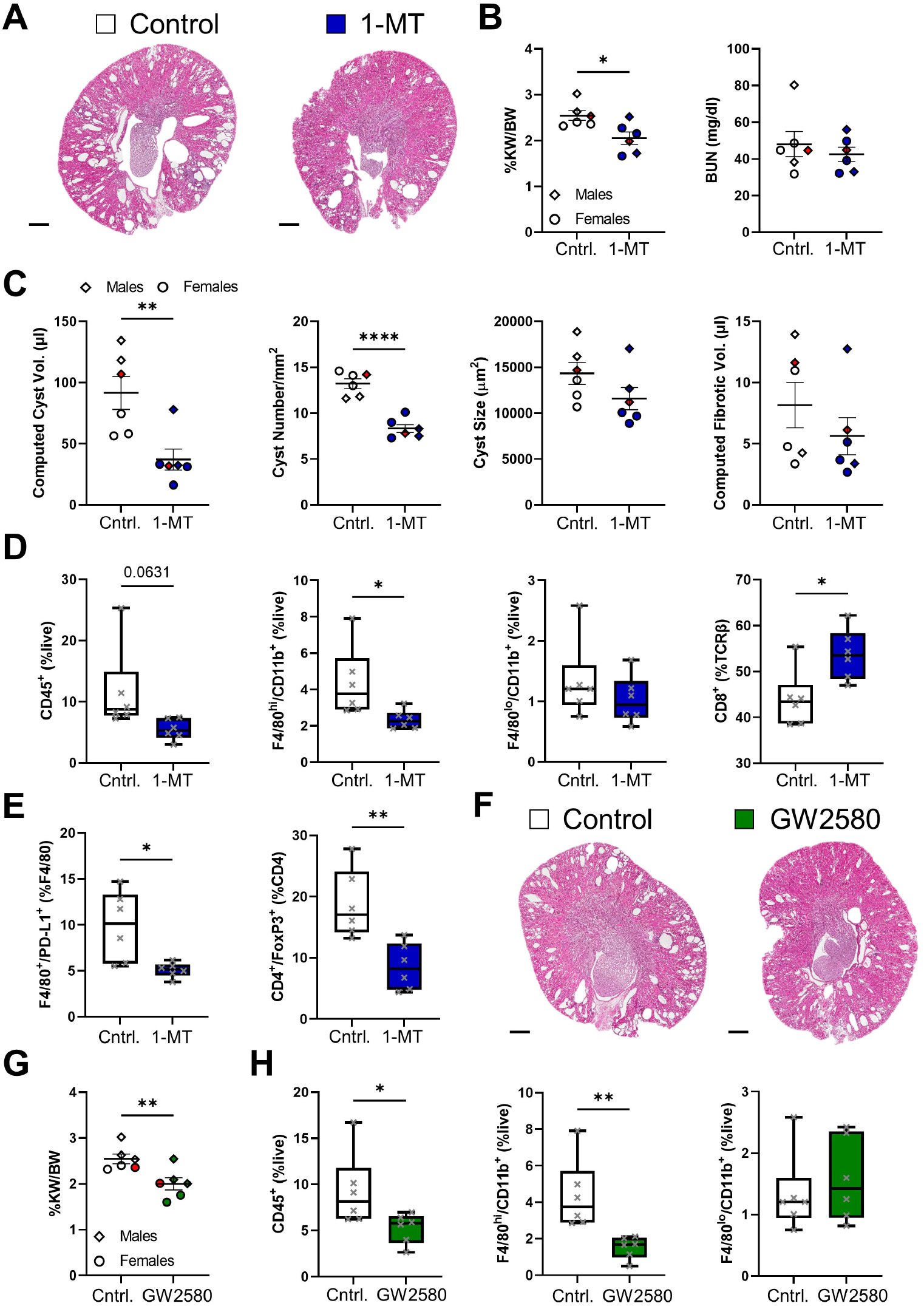
Treatment with a tryptophan analog shows therapeutic efficacy for halting PKD progression in an orthologous ADPKD model. Results obtained from *Pkd1*^RC/RC^ mice treated with (blue) or without (white) 1-MT. (A) H&E cross sections. Quantification of (B) %KW/BW and BUN, (C) cystic and fibrotic volume (cyst/fibrotic index [Supplemental Figure 5] multiplied by KW), cyst number, and cyst size. *Pkd1*^RC/RC^ mice treated with 1-MT show significantly reduced PKD severity compared to control (Cntrl.) BUN or fibrotic volume/index were not changed significantly. (D, E) Quantification of flow cytometry data of the 1-MT intervention experiment. 1-MT treated animals have (D) reduced numbers in overall immune cells (CD45^+^), resident macrophages (F4/80^hi^; CD11b^+^) but not infiltrating macrophages (F4/80^to^; CD11b^+^), and increased numbers of CD8^+^ T cells as percent of all T cells (TCRβ^+^), as well as (E) reduced expression of PD-L1 on macrophages (F4/80^+^) and reduced numbers of TRegs (CD4^+^/FoxP3^+^), both suggesting a less immunosuppressed cystic microenvironment. Results obtained from *Pkd1*^RC/RC^ mice treated with (green) or without (white) the CSF-1R inhibitor GW2580. (F) H&E cross sections and (G) %KW/BW quantification, both showing reduced PKD severity in treated animals. (H) Confirmation of a reduction of resident but not infiltrating macrophages in treated versus control mice and a significant reduction of CD45^+^ cells. Red data points depict the animals shown in (A) or (F) per respective experiment. Animals in the Cntrl. groups served as control for both the 1-MT and the GW2580 treatment. Scale bars: 500μm, Treatment (1-MT and GW258): 4-7weeks of age, N=3males (diamond), 3females (circle)/group. Statistics: Graphs: (B, C, G) mean ± SEM, (D, E, H) box plot, whiskers 10-90^th^ percentile; Analyses: unpaired t test. P *<0.05, **<0.01, ****<0.0001, comparisons with non-significant statistics are not shown.

Immune cell- (CD45^+^), NK cell-, DC-, neutrophil-, and macrophage numbers decreased in 1-MT-treated animals versus controls, although not all declines reached significance (**Figure 4D, Supplemental Figure 5B-D**). Interestingly, upon 1-MT treatment, the number of resident macrophages (F4/80^hi^; CD11b^+^) preferentially decreased, while the number of infiltrating macrophages (F4/80^lo^; CD11b^+^) remained unchanged (**Figure 4D**). 1-MT treatment, similarly to genetic loss of *Ido1*, resulted in a shift of the T cell (TCRβ^+^) population towards CD8^+^ T cells and a decrease of double negative T cell numbers within the kidney; no changes were detected in overall T cell numbers or subtypes (**Figure 4D, Supplemental Figure 5E**). 1-MT treated animals also had significantly reduced expression of PD-L1 on macrophages (F4/80^+^), and a trend towards reduced expression on epithelial cells (EpCAM^+^) (**Figure 4E, Supplemental Figure 5F**). We also observed a significant decrease in TReg (CD4^+^/FoxP3^+^) numbers within the kidneys of 1-MT treated *Pkd1*^RC/RC^ mice compared to control, which reinforces the idea of kynurenines driving TReg differentiation (**Figure 4E**). These data provide preclinical support for IDO1 inhibitors as PKD therapeutic and highlight that pharmaceutical inhibition of IDO1 mimics the immunomodulatory phenotypes observed in the genetic model.

Since we previously established a role of CD8^+^ T cells in the *Pkd1*^RC/RC^ model, we evaluated the role of resident macrophages as a modifier of PKD severity in this model as their numbers preferentially decreased upon IDO1 inhibition(43). We treated 1mo *Pkd1*^RC/RC^ mice with 160mg/kg GW2580, a selective CSF-1R kinase inhibitor, via oral gavage once daily for three weeks. GW2580 has been shown to inhibit PKD progression in the C57Bl/6J CAGG-Cre^ERT2^ *Ift88*^flx/flx^ ischemia-reperfusion-injury model and C57Bl/6J *Cys1*^cpk/cpk^ model(42, 57). *Pkd1*^RC/RC^ mice treated with GW2580 displayed less overall cystic burden as seen by histology and a significant decrease in %KW/BW (**Figure 4F, G**). By flow cytometry, we observed a significant decrease of resident macrophages in GW2580 treated animals versus control while infiltrating macrophage numbers remained unchanged (**Figure 4H**).

## Discussion

Tryptophan and its metabolites regulate a variety of physiological processes including cell growth/maintenance and neuronal function. However, >95% of tryptophan is a substrate for the kynurenine pathway, which controls hyperinflammation and long-term immune tolerance(64, 65). While IDO1 plays a minor role in metabolizing tryptophan to kynurenine under normal conditions, IDO1 levels and IDO1-dependent tryptophan metabolism in macrophages, DCs and epithelial cells is potently induced by inflammatory signals. These include IFNγ, interleukin 6, tumor necrosis factor α, and prostaglandin E2, all of which have been found to be elevated in murine models of PKD or ADPKD patient cyst fluid(43, 65–67). Consistently, we observed increased levels of kynurenines and IDO1 correlative to disease severity and increased IDO1 staining in interstitial cells as well as the cystic epithelium of *Pkd1*^RC/RC^ mice.

Kynurenine and IDO1 levels regulate immunosuppression through suppression of anti-tumorigenic immune cells (DCs, NK-, and effector T cells), expansion of pro-tumorigenic immune cells (TRegs, M2 macrophages, and myeloid-derived suppressor cells), and upregulation of immune checkpoints (PD-1|PD-L1, CTLA4|CD80/86)(60, 68, 69). To date, the functional role of DCs, NK-, and CD4^+^ T cells in PKD progression has not been studied. However, urinary CD4^+^ T cell numbers have been shown to correlate with eGFR decline in ADPKD patients(58). Similarly, kidney M2-like macrophages are known to drive PKD, and we have shown that loss of CD8^+^ T cells enhances cyst growth in the *Pkd1*^RC/RC^ model(43, 51–55, 70–72). Here, we further provide data that kidney TReg numbers are increased in *Pkd1*^RC/RC^ mice compared to wildtype, suggesting they may play a role in PKD. Since genetic loss of *Ido1* in our PKD model not only reduced kidney kynurenine levels back to baseline but also slowed cyst growth, we hypothesized inhibition of immunosuppressive pathways to be an underlying mechanism. In line with this hypothesis, we observed decreased numbers of kidney macrophages and increased numbers of CD8^+^ T cells within the adaptive immune cell population in the kidneys of *Pkd1*^RC/RC^ *Ido1* knockout animals compared to controls. Further, we observed downregulation of PD-1l1PD-L1 expression and a reduction of TReg numbers; all favorable of a CME that supports slowed cyst growth. We also observed fewer DCs, NK cells, and neutrophils in *Pkd1*^RC/RC^ mice when tryptophan metabolism was inhibited, but the functional impact of these microenvironmental changes is less clear. A key limitation of our CME analyses is that we only investigated immune cell numbers and not their function. It is also unclear whether our observed changes in CME composition are due to IDO1 inhibition or due to reduced PKD severity. Future mechanistic studies will need to disjoin these two observations using cell type specific knockouts, delineate what drives IDO1 overexpression in PKD, and outline if the impact of IDO1/kynurenines on immune cell function/composition is cell autonomous or driven by changes within the cystic kidney epithelium.

Beyond immunomodulation, kynurenines may also drive PKD progression through binding to the aryl hydrocarbon receptor, which interacts with a multitude of proteins found to be key modulators of PKD, including mechanistic target of rapamycin, mitogen-activated protein kinases, sirtuin-1, and nuclear factor kappa-light-chain-enhancer of activated B cells(73–77). Interestingly, like annual change in kidney growth and eGFR decline in ADPKD patients, kynurenine levels have also been positively associated with body mass index(10, 78, 79). Further, since tryptophan is an essential amino acid, the impact of various dietary regimens on kynurenine levels has been studied in animal models. Caloric restriction and a ketogenic diet resulted in downregulation of kynurenines, both of which have also been found to slow PKD progression in murine models(16–18, 80, 81).

Together, our data provide convincing evidence that the kynurenine pathway and IDO1 are dysregulated in PKD and that targeting the pathway provides a novel therapeutic platform for disease treatment. It remains unclear why in our genetic model of IDO1 loss, we did not observe a phenotypic difference in PKD severity until 6 months of age compared to control or why some of the observed changes in the CME differed between genetic loss or pharmacological inhibition of IDO1. Regarding the latter, it is interesting that only pharmacological inhibition of IDO1 reduced the number of kidney TRegs, which we hypothesize to be drivers of cyst growth. Similarly, pharmacological inhibition of IDO1 resulted in a selective reduction of resident macrophages and not infiltrating macrophages, whereas genetic loss decreased both populations within the kidney. The data on the contribution of infiltrating macrophages to PKD progression is inconsistent, and a single study suggests that resident macrophages promote cyst growth in a non-orthologous model of PKD(55, 57, 82). The role of these populations has not been studied in the slowly progressive *Pkd1*^RC/RC^ model. While we provide preliminary evidence that inhibition of resident macrophages in this model slows cyst growth, more detailed studies are needed.

In conclusion, our data strongly support the testing of FDA approved IDO1 inhibitors such as epacadostat in long-term preclinical ADPKD studies, with a goal of clinical translation. Similarly, we believe that the impact of dysregulated metabolism on immune cell function in PKD warrants further investigation and could emerge as a promising new therapeutic platform for combination treatment approaches with epithelial-centric drugs such as tolvaptan.

## Concise Methods

Full methods are available in the Supplemental Material.

### Mouse models

The homozygous C57Bl/6J p.R3277C (*Pkd1*^RC/RC^)(43–45) model was crossed with C57Bl/6J *Ido1* knockout mice *(Ido1*^-/-^, Jackson Laboratory, stock #005867) to generate C57Bl/6J *Pkd1*^RC/RC^;*Ido1*^-/-^ and C57Bl/6 *Pkd1*^RC/RC^;*Ido1*^+/+^ animals.

### IDO1 and CSF-1R inhibition

One-month-old C57Bl/6J *Pkd1*^RC/RC^ mice were treated by oral gavage for three weeks: twice daily with 400mg/kg 1-Methyl-D-tryptophan (Sigma-Aldrich, #452483, IDO1 inhibitor), once daily with 160mg/kg GW2580 (LC Laboratories, #G-5903, CSF-1R inhibitor), or once daily with 0.5% hydroxypropyl methyl cellulose, 0.1% Tween 80 (control).

### Human samples

De-identified ADPKD patient cyst cells were obtained from the Baltimore Polycystic Kidney Disease Research and Clinical Core Center (NIDDK, P30DK090868).

### Cell culture

Cell lines (RCTE & 9-12) used have been previously described(83). Cyst cells and cell lines were grown in DMEM/F12 50I50 +L-glutamine, 15nM HEPES (Corning Inc. +10% fetal bovine serum (VWR International) and 1% penicillin-streptomycin (Corning).

### IFNγ stimulation

RCTE and 9-12 cells were treated for 24 hours with recombinant human IFNγ (PeproTech, #300-02) at 100ng/ml or PBS control.

### Kidney function analyses

Blood urea nitrogen was measured following the manufacturer’s protocol (QuantiChrom Urea Assay Kit, BioAssay Systems, # 501078333).

### Histomorphometric analysis

Kidney cystic index, cyst size, and cyst number were analyzed using a custom-built NIS-Elements AR v4.6 macro (Nikon). Fibrotic area was analyzed from picrosirius red-stained kidney sections and visualized using an Olympus BX41 microscope (Olympus Corporation) with a linear polarizer. Images were obtained and quantified as previously described(43).

### Immunofluorescence labeling and quantification

Paraffin embedded tissues were processed and stained as previously described(43). Primary and secondary antibodies: 1° anti-mouse IDO1 (clone M1D0-48; BioLegend, 1:50), 2° AF594 goat anti-rabbit IgG (Life Technologies, 1:1000); 1° anti-mouse E-cadherin (clone 36; BD Transduction Laboratories, 1:100), 2° AF488 goat anti-rat IgG (Life Technologies, 1:1000). Sections were mounted with Vectashield mounting medium with DAPI and visualized with a Nikon *Eclipse Ti* microscope.

### Western blotting

Whole cell lysates from cell lines, cells derived from human cysts, or homogenized mouse tissue were prepared in RIPA cell lysis buffer with protease inhibitor (#P8340; Sigma Aldrich). Protein samples were separated by SDS-PAGE-electrophoresis and transferred to PVDF membranes. Membranes were incubated with primary antibodies overnight and detected using HRP-linked secondary antibodies and ECL detection reagents (1° anti-mouse IDO1 [clone M1D0-48; BioLegend, 1:400], 2° anti-rat-HRP [1:5,000], 1° anti-mouse GAPDH [polyclonal FL-335, Santa Cruz Biotechnology, 1:500], 2° anti-rabbit-HRP [1:5,000]). Densitometry on X-ray films was quantified using ImageJ.

### Single cell suspension

Single cell suspensions were prepared as previously described(43). Kidneys harvested from terminal dissection were mechanically dissociated and digested in DMEM/F12, Liberase TL, and DNaseI at 37°C for 30 minutes. Dissociation was completed using a 18G needle and quenched using FA3 buffer. Cells were passed through a 100μm filter, incubated with red blood cell lysis buffer, and then passed through a 70μm filter. Cells were suspended in FA3 buffer for flow cytometry staining.

### Flow Cytometry

Flow cytometry was performed as previously described(43). Detailed methods, antibodies used, and gating strategies are outlined in the Supplemental Material. The blocked, viability dye-stained, cell suspension was split in two and used for different flow cytometry panels. Panel 1 (innate immune- and epithelial cells): CD45-FITC, CD11c-PE, F4/80-PElDazzle594, CD11b-PerCP-Cy5.5, Gr-1-PElCy7, PD-L1-APC, MHCII-DyLight680, EpCAM-APC-eFluor780, and NKp46-eFluor450. Panel 2 (T cells): CD44-FITC, PD-1-PE, CD45-PE-CF594, TCRβ-PE-Cyanine5, CD69-PE-Cy7, CD8-Alexa Fluor 700, CD4-APC/Cyanine7, Ki-67-APC, and FoxP3-eFluor450. Stained cells were analyzed on the Gallios Flow Cytometer. All data were analyzed using Kaluza Analysis v2.1 (Beckman Coulter).

### Metabolomics - Liquid chromatography tandem mass spectrometry (LC/MS-MS)

<u>Semi-quantitative targeted metabolomics</u> was performed following a validated approach(26, 84). Kidney tissue was homogenized in 80% (v/v) cooled methanol, incubated for protein precipitation, dried in a SpeedVac concentrator centrifuge (Savant, ThermoFisher), and reconstituted in water/methanol (80:20 v/v). Selected multiple reaction monitoring of 250 metabolites using a positive/negative ion-switching high-performance liquid chromatography-tandem mass spectrometry (5500 QTRAP HPLC-MS/MS21) was used for analysis. MultiQuant (v2.1.1, Sciex) software was used for data processing of the 250 unique metabolites.

<u>Kynurenines</u> were analyzed using a modified protocol(85). Frozen tissue was homogenized in 0.5mL formic acid (10% in water)Imethanol (30/70, v/v). The extraction solution was enriched with isotope labeled internal standards and samples were vortexed and centrifuged at 26,000×g for 20min. Supernatant was transferred into HPLC vials with glass inserts. LC-MS/MS was performed on an Agilent Technologies 1200 HPLC system connected to an ABSCIEX 5500 QTRAP mass spectrometer equipped with a turbo ion spray source operated in electrospray mode. LC separation was carried out on an Atlantis T3 3μm (2.1×50mm) column (Waters Corp.) using a mobile phase consisting of 0.1% formic acid in water (Solvent A) and acetonitrile (Solvent B). All analytes were detected in positive ion multiple reaction moTukey’s multiple comparison testnitoring mode.

<u>Data Analysis</u> was performed using MetaboAnalyst v4.0(86).

### Statistical analysis

Data were analyzed using PRISM9 (GraphPad Software). Data are depicted as mean ± SEM or box plot with whiskers of 10-90^th^ percentile; single data points are depicted. Analyses were performed as unpaired t test or one-way ANOVA with Tukey’s multiple comparison test. P-values are denoted by *(P<0.05), **(P<0.01), ***(P<0.001), and ****(P<a0.0001).

### Study Approval

All animal procedures were performed in an AAALAC-accredited facility in accordance with the *Guide for the Care and Use of Laboratory Animals(87)* and approved by the University of Colorado Anschutz Medical Campus Institutional Animal Care and Use Committee (protocol #33, #685).

## Author Contribution

DTN, EKK and KH designed the research study and BYG, MBC, and RAN provide guidance on the study design; DTN, EKK, ND, ETC, JK, and KH conducted the experiments and acquired the data; DTN, JK, and KH analyzed the data; RAN, JK, and KH wrote the manuscript; EKK, ND, BYG, MBC and ETC provide feedback on the manuscript. Authorship order among DTN and EKK, co-first authors, was decided based on overall time and intellectual contribution to the study.

## Acknowledgements

Support for this research was provided by NIH NIDDK K01 DK114164 (KH), NIH NIDDK P30 DK090868 (P&F KH), NIH R01 DK114424 (JK), NIH R01 DK114424-03S1 (JK), NIH NIDDK T32 5T32DK007135 (EKK), the PKD Foundation Research Grant 216G18a (KH), and the Zell Family Foundation (KH, RAN). The *Pkd1*^RC/RC^ mouse model was provided by the Mayo Clinic Robert M. and Billie Kelley Pirnie Translational Polycystic Kidney Disease Research Center (NIDDK P30 DK090728). Human, ADPKD patient, cyst-derived cells were received from the Baltimore Polycystic Kidney Disease Research and Clinical Core Center (NIDDK, P30DK090868). George S. DeBeck (Nikon Instruments Inc.) aided in the development of the histomorphometric analysis macro for the cystic kidney.

## References

1. Bergmann C, Guay-Woodford LM, Harris PC, Horie S, Peters DJM, and Torres VE. Polycystic kidney disease. Nat Rev Dis Primers. 2018;4(1):50.

2. Torres VE, Harris PC, and Pirson Y. Autosomal dominant polycystic kidney disease. Lancet. 2007;369(9569):1287–301.

3. Reule S, Sexton DJ, Solid CA, Chen SC, Collins AJ, and Foley RN. ESRD from autosomal dominant polycystic kidney disease in the United States, 2001-2010. Am J Kidney Dis. 2014;64(4):592–9.

4. System USRD. Bethesda, MD: National Institutes of Health, National Institute of Diabetes and Digestive and Kidney Diseases; 2018.

5. Torres VE, Chapman AB, Devuyst O, Gansevoort RT, Grantham JJ, Higashihara E, et al. Tolvaptan in patients with autosomal dominant polycystic kidney disease. N Engl J Med. 2012;367(25):2407–18.

6. Torres VE, Chapman AB, Devuyst O, Gansevoort RT, Perrone RD, Koch G, et al. Tolvaptan in Later-Stage Autosomal Dominant Polycystic Kidney Disease. N Engl J Med. 2017;377(20):1930–42.

7. Cornec-Le Gall E, Torres VE, and Harris PC. Genetic Complexity of Autosomal Dominant Polycystic Kidney and Liver Diseases. J Am Soc Nephrol. 2018;29(1):13–23.

8. Hopp K, Cornec-Le Gall E, Senum SR, Te Paske I, Raj S, Lavu S, et al. Detection and characterization of mosaicism in autosomal dominant polycystic kidney disease. Kidney Int. 2020;97(2):370–82.

9. Harris PC, Bae KT, Rossetti S, Torres VE, Grantham JJ, Chapman AB, et al. Cyst number but not the rate of cystic growth is associated with the mutated gene in autosomal dominant polycystic kidney disease. J Am Soc Nephrol. 2006;17(11):3013–9.

10. Nowak KL, You Z, Gitomer B, Brosnahan G, Torres VE, Chapman AB, et al. Overweight and Obesity Are Predictors of Progression in Early Autosomal Dominant Polycystic Kidney Disease. J Am Soc Nephrol. 2018;29(2):571–8.

11. Torres VE, Abebe KZ, Schrier RW, Perrone RD, Chapman AB, Yu AS, et al. Dietary salt restriction is beneficial to the management of autosomal dominant polycystic kidney disease. Kidney Int. 2017;91(2):493–500.

12. Takakura A, Contrino L, Zhou X, Bonventre JV, Sun Y, Humphreys BD, et al. Renal injury is a third hit promoting rapid development of adult polycystic kidney disease. Hum Mol Genet. 2009;18(14):2523–31.

13. Nowak KL, and Hopp K. Metabolic Reprogramming in Autosomal Dominant Polycystic Kidney Disease: Evidence and Therapeutic Potential. Clin J Am Soc Nephrol. 2020;15(4):577–84.

14. Menezes LF, and Germino GG. The pathobiology of polycystic kidney disease from a metabolic viewpoint. Nat Rev Nephrol. 2019;15(12):735–49.

15. Padovano V, Podrini C, Boletta A, and Caplan MJ. Metabolism and mitochondria in polycystic kidney disease research and therapy. Nat Rev Nephrol. 2018;14(11):678–87.

16. Warner G, Hein KZ, Nin V, Edwards M, Chini CC, Hopp K, et al. Food Restriction Ameliorates the Development of Polycystic Kidney Disease. J Am Soc Nephrol. 2016;27(5):1437–47.

17. Kipp KR, Rezaei M, Lin L, Dewey EC, and Weimbs T. A mild reduction of food intake slows disease progression in an orthologous mouse model of polycystic kidney disease. Am J Physiol Renal Physiol. 2016;310(8):F726–F31.

18. Torres JA, Kruger SL, Broderick C, Amarlkhagva T, Agrawal S, Dodam JR, et al. Ketosis Ameliorates Renal Cyst Growth in Polycystic Kidney Disease. Cell Metab. 2019;30(6):1007–23 e5.

19. Rowe I, Chiaravalli M, Mannella V, Ulisse V, Quilici G, Pema M, et al. Defective glucose metabolism in polycystic kidney disease identifies a new therapeutic strategy. Nat Med. 2013;19(4):488–93.

20. Chiaravalli M, Rowe I, Mannella V, Quilici G, Canu T, Bianchi V, et al. 2-Deoxy-d-Glucose Ameliorates PKD Progression. J Am Soc Nephrol. 2016;27(7):1958–69.

21. Lakhia R, Yheskel M, Flaten A, Quittner-Strom EB, Holland WL, and Patel V. PPARalpha agonist fenofibrate enhances fatty acid beta-oxidation and attenuates polycystic kidney and liver disease in mice. Am J Physiol Renal Physiol. 2018;314(1):F122–F31.

22. Flowers EM, Sudderth J, Zacharias L, Mernaugh G, Zent R, DeBerardinis RJ, et al. Lkb1 deficiency confers glutamine dependency in polycystic kidney disease. Nat Commun. 2018;9(1):814.

23. Trott JF, Hwang VJ, Ishimaru T, Chmiel KJ, Zhou JX, Shim K, et al. Arginine reprogramming in ADPKD results in arginine-dependent cystogenesis. Am J Physiol Renal Physiol. 2018;315(6):F1855–F68.

24. Ramalingam H, Kashyap S, Cobo-Stark P, Flaten A, Chang CM, Hajarnis S, et al. A methionine-Mettl3-N(6)-methyladenosine axis promotes polycystic kidney disease. Cell Metab. 2021;33(6):1234–47 e7.

25. Grams ME, Tin A, Rebholz CM, Shafi T, Kottgen A, Perrone RD, et al. Metabolomic Alterations Associated with Cause of CKD. Clin J Am Soc Nephrol. 2017;12(11):1787–94.

26. Baliga MM, Klawitter J, Christians U, Hopp K, Chonchol M, Gitomer BY, et al. Metabolic profiling in children and young adults with autosomal dominant polycystic kidney disease. Sci Rep. 2021;11(1):6629.

27. Badawy AA. Kynurenine Pathway of Tryptophan Metabolism: Regulatory and Functional Aspects. Int J Tryptophan Res. 2017;10:1178646917691938.

28. Schmiedel BJ, Singh D, Madrigal A, Valdovino-Gonzalez AG, White BM, Zapardiel-Gonzalo J, et al. Impact of Genetic Polymorphisms on Human Immune Cell Gene Expression. Cell. 2018;175(6):1701–15 e16.

29. Monaco G, Lee B, Xu W, Mustafah S, Hwang YY, Carre C, et al. RNA-Seq Signatures Normalized by mRNA Abundance Allow Absolute Deconvolution of Human Immune Cell Types. Cell Rep. 2019;26(6):1627–40 e7.

30. Debnath S, Velagapudi C, Redus L, Thameem F, Kasinath B, Hura CE, et al. Tryptophan Metabolism in Patients With Chronic Kidney Disease Secondary to Type 2 Diabetes: Relationship to Inflammatory Markers. Int J Tryptophan Res. 2017;10:1178646917694600.

31. Munipally PK, Agraharm SG, Valavala VK, Gundae S, and Turlapati NR. Evaluation of indoleamine 2,3-dioxygenase expression and kynurenine pathway metabolites levels in serum samples of diabetic retinopathy patients. Arch Physiol Biochem. 2011;117(5):254–8.

32. Prendergast GC. Cancer: Why tumours eat tryptophan. Nature. 2011; 478(7368):192–4.

33. Rhee EP, Clish CB, Ghorbani A, Larson MG, Elmariah S, McCabe E, et al. A combined epidemiologic and metabolomic approach improves CKD prediction. J Am Soc Nephrol. 2013;24(8):1330–8.

34. Huang JY, Butler LM, Midttun O, Ulvik A, Wang R, Jin A, et al. A prospective evaluation of serum kynurenine metabolites and risk of pancreatic cancer. PLoS One. 2018;13(5):e0196465.

35. Routy JP, Routy B, Graziani GM, and Mehraj V. The Kynurenine Pathway Is a Double-Edged Sword in Immune-Privileged Sites and in Cancer: Implications for Immunotherapy. Int J Tryptophan Res. 2016;9:67–77.

36. Brochez L, Chevolet I, and Kruse V. The rationale of indoleamine 2,3-dioxygenase inhibition for cancer therapy. Eur J Cancer. 2017;76:167–82.

37. Seeger-Nukpezah T, Geynisman DM, Nikonova AS, Benzing T, and Golemis EA. The hallmarks of cancer: relevance to the pathogenesis of polycystic kidney disease. Nat Rev Nephrol. 2015;11(9):515–34.

38. Liu M, Wang X, Wang L, Ma X, Gong Z, Zhang S, et al. Targeting the IDO1 pathway in cancer: from bench to bedside. J Hematol Oncol. 2018;11(1):100.

39. Bader JE, Voss K, and Rathmell JC. Targeting Metabolism to Improve the Tumor Microenvironment for Cancer Immunotherapy. Mol Cell. 2020;78(6):1019–33.

40. Patel CH, Leone RD, Horton MR, and Powell JD. Targeting metabolism to regulate immune responses in autoimmunity and cancer. Nat Rev Drug Discov. 2019;18(9):669–88.

41. Zimmerman KA, Hopp K, and Mrug M. Role of chemokines, innate and adaptive immunity. Cell Signal. 2020;73:109647.

42. Li Z, Zimmerman KA, and Yoder BK. Resident Macrophages in Cystic Kidney Disease. Kidney360. 2021;2(1):167–75.

43. Kleczko EK, Marsh KH, Tyler LC, Furgeson SB, Bullock BL, Altmann CJ, et al. CD8(+) T cells modulate autosomal dominant polycystic kidney disease progression. Kidney Int. 2018;94(6):1127–40.

44. Arroyo J, Escobar-Zarate D, Wells HH, Constans MM, Thao K, Smith JM, et al. The genetic background significantly impacts the severity of kidney cystic disease in the Pkd1(RC/RC) mouse model of autosomal dominant polycystic kidney disease. Kidney Int. 2021.

45. Hopp K, Ward CJ, Hommerding CJ, Nasr SH, Tuan HF, Gainullin VG, et al. Functional polycystin-1 dosage governs autosomal dominant polycystic kidney disease severity. J Clin Invest. 2012;122(11):4257–73.

46. Wirthgen E, Hoeflich A, Rebl A, and Gunther J. Kynurenic Acid: The Janus-Faced Role of an Immunomodulatory Tryptophan Metabolite and Its Link to Pathological Conditions. Front Immunol. 2017;8:1957.

47. Zhou X, Fan LX, Sweeney WE, Jr., Denu JM, Avner ED, and Li X. Sirtuin 1 inhibition delays cyst formation in autosomal-dominant polycystic kidney disease. J Clin Invest. 2013;123(7):3084–98.

48. Robinson CM, Shirey KA, and Carlin JM. Synergistic transcriptional activation of indoleamine dioxygenase by IFN-gamma and tumor necrosis factor-alpha. J Interferon Cytokine Res. 2003;23(8):413–21.

49. Meireson A, Devos M, and Brochez L. IDO Expression in Cancer: Different Compartment, Different Functionality? Front Immunol. 2020;11:531491.

50. Nemenoff RA, Kleczko EK, and Hopp K. Renal double negative T cells: unconventional cells in search of a function. Ann Transl Med. 2019;7(Suppl 8):S342.

51. Swenson-Fields KI, Vivian CJ, Salah SM, Peda JD, Davis BM, van Rooijen N, et al. Macrophages promote polycystic kidney disease progression. Kidney Int. 2013;83(5):855–64.

52. Karihaloo A, Koraishy F, Huen SC, Lee Y, Merrick D, Caplan MJ, et al. Macrophages promote cyst growth in polycystic kidney disease. J Am Soc Nephrol. 2011;22(10):1809–14.

53. Locatelli L, Cadamuro M, Spirli C, Fiorotto R, Lecchi S, Morell CM, et al. Macrophage recruitment by fibrocystin-defective biliary epithelial cells promotes portal fibrosis in congenital hepatic fibrosis. Hepatology. 2016;63(3):965–82.

54. Viau A, Bienaime F, Lukas K, Todkar AP, Knoll M, Yakulov TA, et al. Cilia-localized LKB1 regulates chemokine signaling, macrophage recruitment, and tissue homeostasis in the kidney. EMBO J. 2018.

55. Cassini MF, Kakade VR, Kurtz E, Sulkowski P, Glazer P, Torres R, et al. Mcp1 Promotes Macrophage-Dependent Cyst Expansion in Autosomal Dominant Polycystic Kidney Disease. J Am Soc Nephrol. 2018;29(10):2471–81.

56. Zimmerman KA, Huang J, He L, Revell DZ, Li Z, Hsu JS, et al. Interferon Regulatory Factor-5 in Resident Macrophage Promotes Polycystic Kidney Disease. Kidney360. 2020;1(3):179–90.

57. Zimmerman KA, Song CJ, Li Z, Lever JM, Crossman DK, Rains A, et al. Tissue-Resident Macrophages Promote Renal Cystic Disease. J Am Soc Nephrol. 2019;30(10):1841–56.

58. Zimmerman KA, Gonzalez NM, Chumley P, Chacana T, Harrington LE, Yoder BK, et al. Urinary T cells correlate with rate of renal function loss in autosomal dominant polycystic kidney disease. Physiol Rep. 2019;7(1):e13951.

59. Mezrich JD, Fechner JH, Zhang X, Johnson BP, Burlingham WJ, and Bradfield CA. An interaction between kynurenine and the aryl hydrocarbon receptor can generate regulatory T cells. J Immunol. 2010;185(6):3190–8.

60. Liu Y, Liang X, Dong W, Fang Y, Lv J, Zhang T, et al. Tumor-Repopulating Cells Induce PD-1 Expression in CD8(+) T Cells by Transferring Kynurenine and AhR Activation. Cancer Cell. 2018;33(3):480–94 e7.

61. Nguyen D, Kleczko EK, Gitomer B, Clambey E, Chonchol M, Klawitter J, et al. Targeting Immunosuppressive Pathways Reduces ADPKD Progression in a Relevant Murine Model. J Am Soc Nephrol. 2019;30(SA-PO461):882.

62. Qian F, Liao J, Villella J, Edwards R, Kalinski P, Lele S, et al. Effects of 1-methyltryptophan stereoisomers on IDO2 enzyme activity and IDO2-mediated arrest of human T cell proliferation. Cancer Immunol Immunother. 2012;61(11):2013–20.

63. Gunther J, Dabritz J, and Wirthgen E. Limitations and Off-Target Effects of Tryptophan-Related IDO Inhibitors in Cancer Treatment. Front Immunol. 2019;10:1801.

64. Platten M, Nollen EAA, Rohrig UF, Fallarino F, and Opitz CA. Tryptophan metabolism as a common therapeutic target in cancer, neurodegeneration and beyond. Nat Rev Drug Discov. 2019;18(5):379–401.

65. Sorgdrager FJH, Naude PJW, Kema IP, Nollen EA, and Deyn PP. Tryptophan Metabolism in Inflammaging: From Biomarker to Therapeutic Target. Front Immunol. 2019;10:2565.

66. Munn DH. Indoleamine 2,3-dioxygenase, tumor-induced tolerance and counter-regulation. Curr Opin Immunol. 2006;18(2):220–5.

67. Zhai L, Spranger S, Binder DC, Gritsina G, Lauing KL, Giles FJ, et al. Molecular Pathways: Targeting IDO1 and Other Tryptophan Dioxygenases for Cancer Immunotherapy. Clin Cancer Res. 2015;21(24):5427–33.

68. Wang XF, Wang HS, Wang H, Zhang F, Wang KF, Guo Q, et al. The role of indoleamine 2,3-dioxygenase (IDO) in immune tolerance: focus on macrophage polarization of THP-1 cells. Cell Immunol. 2014;289(1-2):42–8.

69. Hornyak L, Dobos N, Koncz G, Karanyi Z, Pall D, Szabo Z, et al. The Role of Indoleamine-2,3-Dioxygenase in Cancer Development, Diagnostics, and Therapy. Front Immunol. 2018;9:151.

70. Mrug M, Zhou J, Woo Y, Cui X, Szalai AJ, Novak J, et al. Overexpression of innate immune response genes in a model of recessive polycystic kidney disease. Kidney Int. 2008;73(1):63–76.

71. Zimmerman KA, Song CJ, Li Z, Lever JM, Crossman DK, Rains A, et al. Tissue-Resident Macrophages Promote Renal Cystic Disease. J Am Soc Nephrol. 2019.

72. Zimmerman KA, Huang J, He L, Revell DZ, Li Z, Hsu J-S, et al. Interferon Regulatory Factor 5 in Resident Macrophage Promotes Polycystic Kidney Disease. Kidney 360. 2020:10.34067/KID.0001052019.

73. Diry M, Tomkiewicz C, Koehle C, Coumoul X, Bock KW, Barouki R, et al. Activation of the dioxin/aryl hydrocarbon receptor (AhR) modulates cell plasticity through a JNK-dependent mechanism. Oncogene. 2006;25(40):5570–4.

74. Bessede A, Gargaro M, Pallotta MT, Matino D, Servillo G, Brunacci C, et al. Aryl hydrocarbon receptor control of a disease tolerance defence pathway. Nature. 2014;511(7508):184–90.

75. Denison MS, and Nagy SR. Activation of the aryl hydrocarbon receptor by structurally diverse exogenous and endogenous chemicals. Annu Rev Pharmacol Toxicol. 2003;43:309–34.

76. Tan Z, Chang X, Puga A, and Xia Y. Activation of mitogen-activated protein kinases (MAPKs) by aromatic hydrocarbons: role in the regulation of aryl hydrocarbon receptor (AHR) function. Biochem Pharmacol. 2002;64(5-6):771–80.

77. Saigusa T, and Bell PD. Molecular pathways and therapies in autosomal-dominant polycystic kidney disease. Physiology (Bethesda). 2015;30(3):195–207.

78. Favennec M, Hennart B, Caiazzo R, Leloire A, Yengo L, Verbanck M, et al. The kynurenine pathway is activated in human obesity and shifted toward kynurenine monooxygenase activation. Obesity (Silver Spring). 2015;23(10):2066–74.

79. Nowak KL, Steele C, Gitomer B, Wang W, Ouyang J, and Chonchol MB. Overweight and Obesity and Progression of ADPKD. Clin J Am Soc Nephrol. 2021;16(6):908–15.

80. Heischmann S, Gano LB, Quinn K, Liang LP, Klepacki J, Christians U, et al. Regulation of kynurenine metabolism by a ketogenic diet. J Lipid Res. 2018;59(6):958–66.

81. Strasser B, Berger K, and Fuchs D. Effects of a caloric restriction weight loss diet on tryptophan metabolism and inflammatory biomarkers in overweight adults. Eur J Nutr. 2015;54(1):101–7.

82. Salah SM, Meisenheimer JD, Rao R, Peda JD, Wallace DP, Foster D, et al. MCP-1 promotes detrimental cardiac physiology, pulmonary edema, and death in the cpk model of polycystic kidney disease. Am J Physiol Renal Physiol. 2019;317(2):F343–F60.

83. Gainullin VG, Hopp K, Ward CJ, Hommerding CJ, and Harris PC. Polycystin-1 maturation requires polycystin-2 in a dose-dependent manner. J Clin Invest. 2015;125(2):607–20.

84. Yuan M, Breitkopf SB, Yang X, and Asara JM. A positive/negative ion-switching, targeted mass spectrometry-based metabolomics platform for bodily fluids, cells, and fresh and fixed tissue. Nat Protoc. 2012;7(5):872–81.

85. Zhu W, Stevens AP, Dettmer K, Gottfried E, Hoves S, Kreutz M, et al. Quantitative profiling of tryptophan metabolites in serum, urine, and cell culture supernatants by liquid chromatography-tandem mass spectrometry. Anal Bioanal Chem. 2011;401(10):3249–61.

86. Chong J, Soufan O, Li C, Caraus I, Li S, Bourque G, et al. MetaboAnalyst 4.0: towards more transparent and integrative metabolomics analysis. Nucleic Acids Res. 2018;46(W1):W486–W94.

87. Research IoLA. Guide for the Care and Use of Laboratory Animals. Washington (DC): National Academies Press; 2011.

