## Supplemental Material for "The Tryptophan Metabolizing Enzyme Indoleamine 2,3-Dioxygenase 1 Regulates Polycystic Kidney Disease Progression"

**Supplemental Methods**

*Experimental Study*

All animal procedures were performed in an AAALAC-accredited facility in accordance with the *Guide for the Care and Use of Laboratory Animals*(1) and approved by the University of Colorado Anschutz Medical Campus Institutional Animal Care and Use Committee (protocol #33, #685). From birth until 24 days of age mice were housed with their parents. At 24 days of age mice were weaned with same sex/age mice which were part of the same study: 3-5 mice/cage. For the rest of the duration of the study mice were housed in the vivarium, which maintains a temperature of ~72ºF, humidity of ~37%, and light cycle of 8pm off, 6am on. All cages were sterile, and animals received hyperchlorinated reverse osmosis water delivered via an automatic watering system and irradiated diet (ENVIGO #2920).

*Mouse models*

Fully inbred, homozygous C57Bl/6J *Pkd1*$^{RC/RC}$ mice were obtained from the Mayo Clinic in 2015 with an approved MTA and were maintained in homozygosity by the study PI, Katharina Hopp(2, 3). Homozygous C57Bl/6J *Pkd1*$^{RC/RC}$ mice were outcrossed every 10$^{th}$ generation to wildtype C57Bl/6J mice obtained from *The Jackson Laboratory* (stock #000664). The C57Bl/6J Ido1 knock-out (*Ido1*$^{-/-}$, stock #005867) mice were purchased from *The Jackson Laboratory*. For the treatment studies all C57Bl/6J *Pkd1*$^{RC/RC}$ experimental animals were from an 8$^{th}$ generation backcross and their genotype was confirmed as previously published(3, 4). For the genetic studies, C57Bl/6J *Pkd1*$^{RC/RC}$ and *Ido1*$^{-/-}$ mice were crossed for two generations. All experimental animals (C57Bl/6J *Pkd1*$^{RC/RC}$ *Ido1*$^{+/+}$ *or* C57Bl/6J *Pkd1*$^{RC/RC}$ *Ido1*$^{+/+}$) were F3 animals. All mice were sacrificed at the indicated 3-month and 6-month time points and both sexes, males and females, were utilized for the study.

*IDO1 and CSF-1R inhibition*

One month old C57Bl/6J *Pkd1*$^{RC/RC}$ mice were treated by oral gavage for three weeks. The following three groups were treated simultaneously (all animals in this study were 8$^{th}$ generation C57Bl/6J *Pkd1*$^{RC/RC}$ mice): IDO1 inhibitor: twice daily, 400 mg/kg, 1-Methyl-D-tryptophan (Sigma-Aldrich, #452483), 80mg/ml dissolved in 0.5% hydroxypropyl methyl cellulose, 0.1% Tween 80; CSF-1R inhibitor: once daily, 160 mg/kg, GW2580 (LC Laboratories, #G-5903), 32mg/ml dissolved in 0.5% hydroxypropyl methyl cellulose, 0.1% Tween 80; Control: once daily, 0.5% hydroxypropyl methyl cellulose, 0.1% Tween 80.

*Human samples*

De-identified ADPKD patient cyst cells were obtained from the Baltimore Polycystic Kidney Disease Research and Clinical Core Center (NIDDK, P30DK090868, contact: Dr. Terry Watnick). Each cell colony was established from a single cyst, was passaged once, and then shipped. For some cysts, multiple primary cell cultures were established.

*Mouse tissue harvest/analysis*

Animals were euthanized by isoflurane exposure followed by cervical dislocation, and the body weight of each animal was recorded. A terminal blood collection was performed by cardiac puncture. The kidneys were harvested and weighed. The left kidney was used for the single cell suspension/flow cytometry. The right kidney was cut into 2-5mm sections at each pole and the center, fixed in 4% paraformaldehyde, and embedded in paraffin for histological analyses.

*Cell Culture*

Cell lines (RCTE & 9-12) used have been previously described(5). RCTE cells are *PKD1*$^{+/+}$ immortalized human renal cortical tubular epithelial cells, and 9-12 cells are *PKD1*$^{-/-}$ immortalized cells derived from human ADPKD cystic epithelium. All primary cyst cells and cell lines were grown in Dulbecco's modified Eagle's medium/Ham's F-12 50/50 mix with L-glutamine and 15nM HEPES (DMEM/F12; Corning; RCTE, 9-12) supplemented with 10% fetal bovine serum (Sigma-Aldrich) and 1% penicillin-streptomycin (Corning). Cells were grown in a humidified incubator with 5% $CO_2$ at 37°C.

*IFN-γ stimulation*

Cultured RCTE and 9-12 cells were grown to 70% confluence on 10cm culture plates. Human recombinant IFN-γ (PeproTech, Rocky Hill, NJ, #300-02) was added to culture medium at 100 ng/ml for 24 hours. Control cells received phosphate-buffered saline. After 24h, cells were harvested and lysed by scraping down plates with ice cold PBS, centrifuging at 300g for 5 minutes, and incubating the cell pellet in RIPA lysis Buffer with 1:100 Protease Inhibitor Cocktail (Sigma-Aldrich, #P8340) for 10 minutes after vortexing. The cells were then centrifuged at 13,000rpm for 5 minutes, and cell lysate supernatant was transferred to clean tubes.

*Kidney Function Analyses*

BUN was measured using the QuantiChrom Urea Assay Kit (BioAssay Systems, # 501078333) according to the manufacturer's protocol. Whole blood was collected via cardiac puncture at time of dissection and collected in heparin-coated tubes. Plasma was isolated via centrifugation. 5 μL of plasma were analyzed per sample and all animals were analyzed in duplicates.

*Histomorphometric analysis*

Image acquisition and analysis was performed as previously described(4). Cystic index, cyst size, and number were analyzed using a custom-built NIS-Elements AR v4.6 macro (Nikon) using three cross-sections per kidney (two poles, and center/pelvis). A cyst was defined as having a minimum feret diameter of 50μm. The cystic index was defined as the percentage of cystic area per kidney cross section, and cyst number was normalized to area. Fibrotic area was analyzed from picrosirius red stained kidney sections and, visualized using an Olympus BX41 microscope with a linear polarizer. Ten random cortical 40x images were analyzed per animal and the percent fibrotic area was calculated from total kidney tissue area. Fibrillar collagen (birefringent area) was quantified using ImageJ. Computed volumes were calculated by multiplying the obtained indices (cystic or fibrotic) with the kidney weight of the same animal.

*Immunofluorescence labeling/quantification*

Tissues were prepared for immunofluorescence labeling as previously described(4). 4μm sections underwent autofluorescence quenching with 0.1% Sudan Black in 70% ethanol, 2h citrate antigen retrieval at 100ºC, and washes in TBST. Primary antibodies used were anti-mouse IDO1 (clone M1D0-48; BioLegend) at 1:50 and anti-mouse E-cadherin (clone 36; BD Transduction laboratories) at 1:100. Secondary antibodies used were AF488 goat anti-rat IgG, AF594 goat anti-rabbit IgG, and AF647/AF488 goat anti-mouse IgG2a (Life Technologies) at 1:1000. Slides were mounted using VectraShield (Vector Laboratories, #H-1200). Image visualization was performed using a Nikon *Eclipse Ti* microscope with a Zyla 4.2sCMOS camera. Image analysis was done using NIS-Elements AR v4.6 (Nikon).

*Western blotting*

Kidney tissue was homogenized in lysis buffer containing RIPA buffer and protease inhibitor (#P8340; Sigma Aldrich) using a Qiagen TissueLyser LT homogenizer. Cells were lysed using RIPA Lysis Buffer with Protease Inhibitor Cocktail. Protein concentration was measured using Protein assay dye reagent concentrate (#5000006, Bio-Rad) and 30μg of samples were loaded onto 4-12% acrylamide gels and ran for 1 hour at 200 V. Cultured cell lysates and human cyst lysates were loaded at 30μg protein per well. Gels were transferred to PVDF membranes at 400 mAmps for 3 hours followed by blocking in 5% BSA. The primary antibodies used were anti-mouse IDO1 (clone mIDO-48, 1:400, BioLegend) and anti-mouse GAPDH (polyclonal FL-335,

1:500, Santa Cruz Biotechnology). Respective secondary anti-rat-HRP (1:5000) and secondary anti-rabbit-HRP (1:5000, Jackson ImmunoResearch, West Grove, PA) antibodies were used and blots were developed using Western Lightning Plus-ECL substrate (#NEL104001EA PerkinElmer, Waltham, MA). Band density of each blot was quantified using Image J software.

*Single cell suspension*

Dissected kidney tissue was mechanically dissociated using razor blades and placed in 3.6mL of DMEM/F12 media (Corning) with 0.4mL Liberase TL (2mg/mL in DMEM/F12, Sigma-Aldrich, #05401020001), and 20µL DNaseI (20K U/mL in Hank's Buffer; Sigma-Aldrich, #D5025). Tissue suspensions were placed in a 37°C shaking water bath for 30 minutes, and the samples were mixed every 10 minutes. To ensure dissociation, the digestion mix was passed through an 18G needle several times. An equal volume of FA3 Buffer (PBS, 10mM HEPES [Corning], 2nM EDTA, 1% FBS [Sigma-Aldrich]) was added to the tissue digestion mixture and strained through a 100 µm filter. Single cells were spun down and washed in FA3 Buffer. The cell pellet was then resuspended in 2 mL of Red Blood Cell (RBC) Lysis Buffer (0.015M $NH_4Cl$, 10mM $KHCO_3$, 0.1mM $Na_2EDTA$, pH 7.2) for exactly 3 minutes at room temperature. The lysis was quenched by adding 13mL of FA3 Buffer. After centrifuging, the cell pellet was resuspended in 10 mL of FA3 Buffer and passed through a 70 µm filter. The resulting pelleted cells were then ready for flow cytometry staining.

*Flow Cytometry*

The single cell suspension was blocked in anti-mouse CD16/CD32 (clone 93; eBioscience, #14-0161-86) at 1:200 on a rocker for 15 min at 4°C. Following, viability dye (LIVE/DEAD Fixable Aqua Dead Cell Stain Kit, Invitrogen, #L34966) was added and the suspension was stained for 15 min at 4 °C. The cells were then washed once with FA3 and then split in half for staining of two different flow cytometry panels.

Panel 1

To one half of the single cell suspension the following conjugated antibodies to surface markers were added: CD45-FITC (clone 30-F11; 1:100; BioLegend), CD11c-PE (clone HL3; 1:100; BD Biosciences), F4/80-PE/Dazzle594 (clone BM8; 1:100; BioLegend), CD11b-PerCP-Cy5.5 (clone M1/70; 1:100; BD Biosciences), Gr-1-PE/Cy7 (clone RB6-8C5; 1:100; BioLegend), PD-L1-APC (clone MIH5; 1:100; eBioscience), MHCII-DyLight680 (clone M5/114.15.2; 1:100; Novus), EpCAM-APC-eFluor780 (clone G8.8; 1:100; eBioscience), NKp46-eFluor450 (clone 29A1.4; 1:100; eBioscience), and CD4-V500 (clone RM4-5; 1:100; BD Biosciences; used only for compensation). Cells were incubated with all antibodies in the dark on a rocker at 4°C for 60 min followed by two washes in FA3 buffer. Finally, cells were resuspended in FA3 Buffer and ran on the Gallios Flow Cytometer Machine (Beckman Coulter). For compensation, single-stained beads (VersaComp Antibody Capture Bead Kit; Beckman Coulter) and a cell-mix of all samples were used.

Panel 2

To the second half of the single cell suspension the following conjugated antibodies were added on day 1: CD44-FITC (clone IM7; 1:100; eBioscience), PD-1-PE (clone RMP1-30; 1:100; BD Biosciences), CD45-PE-CF594 (clone 30-F11; 1:100; eBioscience), TCRβ-PE-Cyanine5 (clone H57-597; 1:100; eBioscience), CD69-PE-Cy7 (clone H1.2F3; 1:100; eBioscience), CD8-Alexa Fluor 700 (clone 53-6.7; 1:100; eBioscience), CD4-APC/Cyanine7 (clone GK1.5; 1:100; BioLegend), and CD4-V500 (clone RM4-5; 1:100; BD Biosciences; used only for compensation). Cells were incubated with the antibody mix in the dark on a rocker at 4°C for 60 min. Following, cells were washed with and incubated in fixation/permeabilization buffer overnight according to the Foxp3/Transcription Factor Staining Buffer Set (eBioscience, #00-5523-00). The next day, conjugated antibodies with intracellular targets were added to cells in permeabilization buffer (Ki-67-APC [clone SolA15; 1:400; eBioscience] and FoxP3-eFluor450 [clone FJK-16s; 1:200;

eBioscience]) and cells were incubated with the antibody mix for 120 minutes on a rocker at 4°C in the dark. Cells were washed twice, resuspended in FA3 buffer and ran on the Gallios Flow Cytometer Machine (Beckman Coulter). Compensation using single stained bead and pooled cell mix was performed.

Data analysis

Flow Cytometry data were analyzed using the Kaluza Analysis v2.1 software (Beckman Coulter). First, compensation for each channel was performed using single-stained beads and confirmed using single-stained pooled cell mix. All samples were then analyzed using the gated workflow shown in **Supplemental Figure 6** in a blinded manner.

*Metabolomics - Liquid chromatography tandem mass spectrometry (LC/MS-MS)*

Semi-quantitative targeted metabolomics

Sample analysis was performed based on a validated approach(6, 7). Kidneys were perfused with ice cold PBS/heparin, and kidneys were dissected and snap frozen in liquid nitrogen. Following, kidney tissue samples (~50-100 mg) were homogenized in adequate volume of 80% (v/v) cooled methanol, incubated for protein precipitation, dried in a SpeedVac concentrator centrifuge (Savant, ThermoFisher, Waltham, MA), and reconstituted in water/methanol. 8 µL of sample was injected onto an Amide XBridge HPLC column (3.5 µm; 4.6 mm inner diameter (i.d.) × 100 mm length; Waters). The mobile phases consisted of HPLC buffer A (pH = 9.0: 95% (vol/vol) water, 5% (vol/vol) acetonitrile, 20 mM ammonium hydroxide, 20 mM ammonium acetate) and HPLC buffer B (100% acetonitrile). The HPLC settings were as follows: from 0 to 3 minutes, the mobile phase was kept at 85% B; from 3 to 22 minutes, the percentage of solvent B was decreased from 85% to 2% and was kept at 2% for additional 3 minutes. At minute 26, solvent B was increased again back to 85% and the column flushed for additional 7 minutes at 85% solvent B.

The Q1 (precursor ion) and Q3 (fragment ion) transitions, the metabolite names, dwell times and the appropriate collision energies (CEs) for both positive and negative ion modes were adapted from(6) with several additional transitions. Q1 and Q3 transitions were set to unit resolution for optimal metabolite ion isolation and selectivity. In addition, the polarity switching (settling) time was set to 50 ms; in 1.42 s using a 3-ms dwell time, we were able to obtain 6-14 scans per metabolite peak. The source temperature was set at 500°C, curtain gas (CUR, nitrogen) at 20, collision gas (CAD, nitrogen) at high, ion source gases 1 and 2 at 33, declustering potential (DP) at +93/-93, entrance potential (EP) at +10/-10, and collision cell exit potential (CXP) at +10/-10 for positive and negative ion modes, respectively. Positive identification of the metabolites of interest was performed through injection of pure compound standards onto the above-described LC-MS/MS platform (confirmation of the fragmentation pattern (MS/MS) and retention time).

Kynurenines

Kynurenines were analyzed using a modification of(8). Briefly, frozen tissue (~15 mg) was weighed and homogenized in 0.5 mL formic acid (10% in water) / methanol (30/70, v/v) using an electric homogenizer. The extraction solution was enriched with isotope labeled internal standard mix (at 10 ng/mL final concentration, see below). Samples were vortexed and centrifuged at 26,000xg for 20 minutes after which the supernatant was transferred into HPLC vials with glass inserts. Calibrator standards and quality control samples were prepared in 0.1% formic acid in water as surrogate matrix. LC-MS/MS was performed on an Agilent Technologies (Santa Clara, CA) 1200 HPLC system connected to an ABSCIEX (Foster City, CA) 5500 QTRAP mass spectrometer equipped with a turbo ion spray source operated in electrospray mode. LC separation was carried out on an Atlantis T3 3µm (2.1x50 mm) column (Waters Corp., Milford, MA) using a mobile phase consisting of 0.1% formic acid in water (Solvent A) and acetonitrile (Solvent B). All analytes were detected in positive ion multiple reaction monitoring (MRM) mode. The following quantifier ion-transitions were monitored TRP (205>118), KYN (209>192), KYNA

(190>144), 3OH KYN (225>208), anthranilic acid (AA) (138>120), picolinic acid (PA) (124>78), QA (168>78), 3-OH AA (154>80), and serotonin (SER) (177>115). The following isotope labeled internal standards have been used: $d_5$-TRP (210.2>147.2), $^{13}C_3$,$^{15}N$-3OH KYN (229.2>110.2), $d_5$-KYNA (195.2>149.2), $^{13}C_4$,$^{15}N$-QA (173>81.2), $^{13}C_6$-KYN (215.2>152.2), $^{13}C^6$-AA (144>98.2), $d_4$-PA (128>82.2), $d_4$-SER (181>164).

Data Analysis

MetaboAnalyst 4.0 (University of Alberta, Canada) was used for statistical analysis of metabolomics data(9). Relative peak intensities were initially normalized to the deuterated internal standards followed by the sum of all integrals and tissue weight. After that, data were log transformed and then Paretto-scaled (mean centered and divided by the square root of the SD of each variable). ANOVA with post-hoc Tukey HSD was used to compare group differences. Analysis of changes in metabolites between different animal groups was performed by utilizing Partial Least Squares-Discriminant Analysis (PLS-DA). False discovery rate (FDR) correction was applied to correct for multiple comparisons (FDR < 0.05 for statistical significance).

*Statistical analysis*

All analyses were performed using PRISM9 (Graphpad Software). Data are depicted as mean ± SEM or box plot with whiskers of 10-90$^{th}$ percentile; single data points are depicted in all instances. Analyses were performed as unpaired t test, or one-way ANOVA with Tukey's multiple comparison test, deepening on data type and group number. P-values are denoted by *(P<0.05), **(P<0.01), ***(P<0.001), and ****(P<0.0001).

### Supplemental Figures

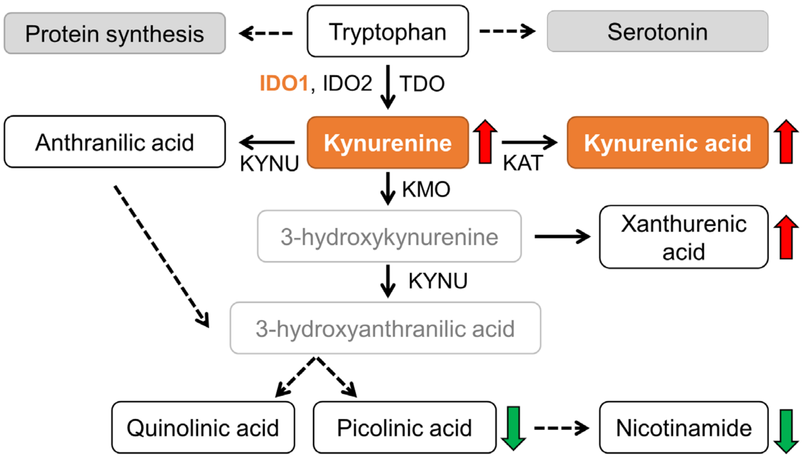


**Supplemental Figure 1 | Schematic of tryptophan metabolism.** Key catabolites analyzed by mass spectrometry are depicted (black/orange; Orange indicates catabolites or enzymes known to have immunomodulatory roles. Colored arrows indicate trend of levels in C57Bl/6J *Pkd1*$^{RC/RC}$ kidneys versus wildtype (red: increased, green: decreased). IDO: indoleamine 2,3-dioxygenase; TDO: tryptophan 2,3-dioxygenase; KATs: kynurenine amino transferases; KMO: kynurenine 3-monooxygenase; KYNU: kynureninase.

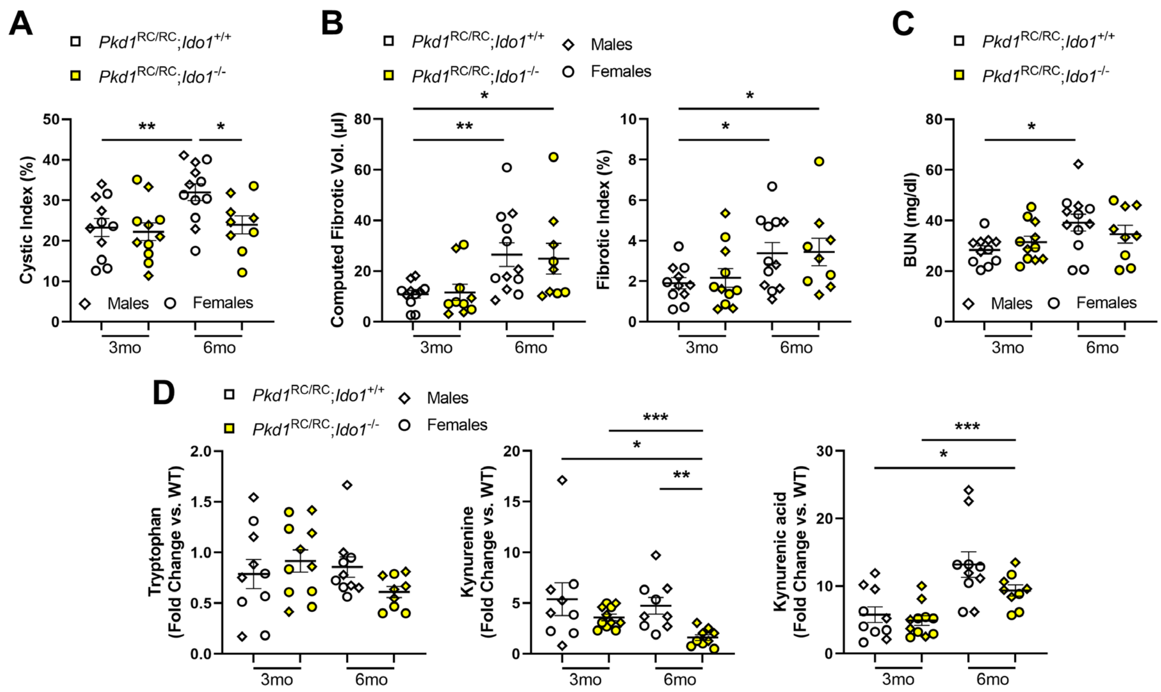


**Supplemental Figure 2 | PKD-related histopathological quantification and metabolomics highlight reduced PKD severity and correction of dysregulated tryptophan metabolism in PKD mice null for *Ido1* versus control.** Quantification of (A) cystic index (B) fibrosis and (C) BUN highlight reduction in cystic kidney disease severity but not fibrotic burden or kidney function decline in *Pkd1*$^{RC/RC}$; *Ido1*$^{+/+}$ (white) versus *Pkd1*$^{RC/RC}$; *Ido1*$^{-/-}$ (yellow) animals. (D) Levels of tryptophan metabolites of kidneys from *Pkd1*$^{RC/RC}$; *Ido1*$^{+/+}$ (white) and *Pkd1*$^{RC/RC}$; *Ido1*$^{-/-}$ (yellow) animals quantified as fold-change compared to genotype, age, and gender matched wildtype (WT) mice. At 3mo and 6mo of age, levels of the immunosuppressive tryptophan metabolite kynurenine are significantly reduced in PKD *Ido1* null animals compared to control, reaching close to WT levels. At 6mo of age, the time point at which a significant reduction in PKD severity is observed in PKD *Ido1* null animals versus control, levels of kynurenic acid also declined, although not significant. N= 5males (diamond), 4-7females (circle) per genotype and time point. Statistics: Graphs: mean ± SEM; Analyses: one-way ANOVA with Tukey's multiple comparison test. P *<0.05, **<0.01, ***<0.001, comparisons with non-significant statistics are not shown.

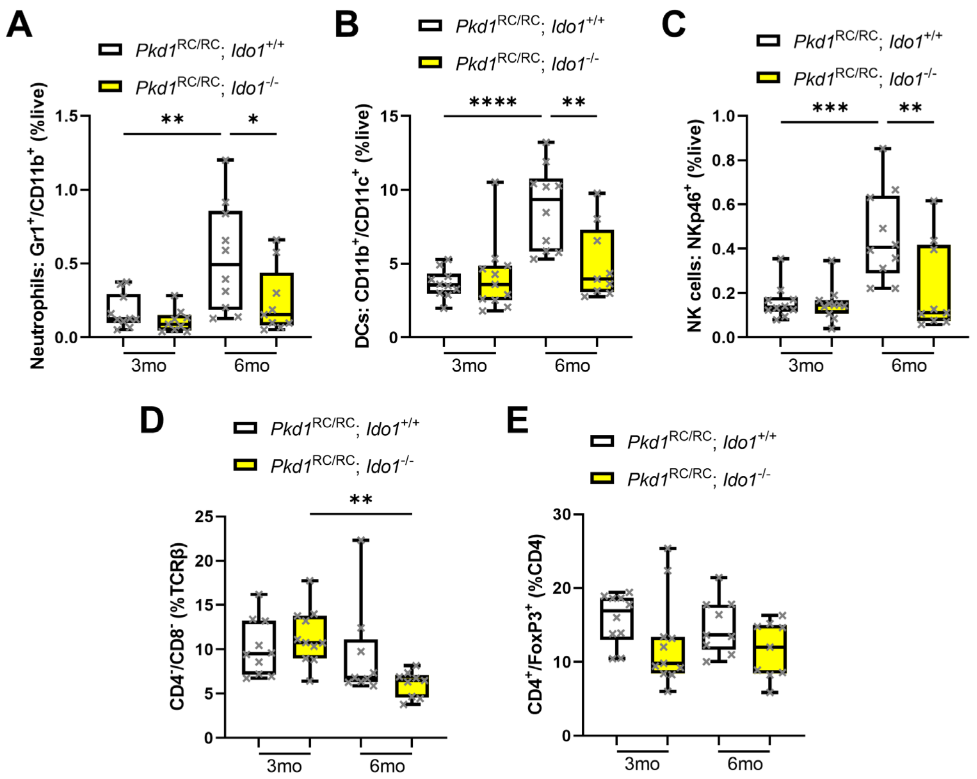


**Supplemental Figure 3 | Kidney innate and adaptive immune cell numbers are altered in PKD *Ido1* null mice compared to control**. Quantification of flow cytometry data obtained from kidney single cell suspensions of *Pkd1*$^{RC/RC}$; *Ido1*$^{+/+}$ (white) and *Pkd1*$^{RC/RC}$; *Ido1*$^{-/-}$ (yellow) animals highlights significantly reduced numbers of (A) neutrophils, (B) dendritic cells, and (C) natural killer cells at 6mo of age, the time point at which reduced PKD severity was observed in animals null for *Ido1*. Further, numbers of (D) double negative T cells as percent of all T cells (TCRβ$^{+}$) were significantly reduced at 6mo and (E) $T_{Regs}$ trend towards a reduction, although not significant, in PKD animals null for *Ido1* versus control at 6mo of age. Statistics: Graphs: box plot, whiskers 10-90$^{th}$ percentile; Analyses: one-way ANOVA with Tukey's multiple comparison test. P *<0.05, **<0.01, ***<0.001, ****<0.0001, comparisons with non-significant statistics are not shown. N= 5males, 4-7females per genotype and time point.

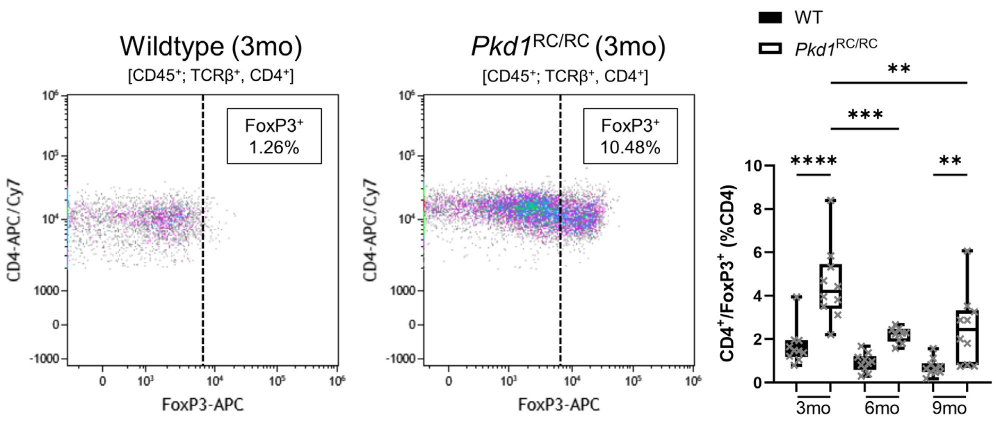


**Supplemental Figure 4 | Kidney regulatory T cell numbers increase in *Pkd1*$^{RC/RC}$ mice compared to wildtype.** Representative flow cytometry plot indicating the gating strategy of regulatory T cells (CD4$^{+}$/Foxp3$^{+}$) in kidney single cell suspensions from C57Bl/6 *Pkd1*$^{RC/RC}$ mice and strain, age, and gender matched wildtype (WT) mice (left). Quantification at 3mo, 6mo, and 9mo (right). Numbers of $T_{Regs}$ are increased at all investigated time points, however the increase is most prominent at milder disease stages (3mo). Statistics: Graphs: box plot, whiskers 10-90$^{th}$ percentile; Analyses: one-way ANOVA with Tukey's multiple comparison test. **<0.01, ***<0.001, ****<0.0001, comparisons with non-significant statistics are not shown. N= 5males, 5females per genotype and time point.

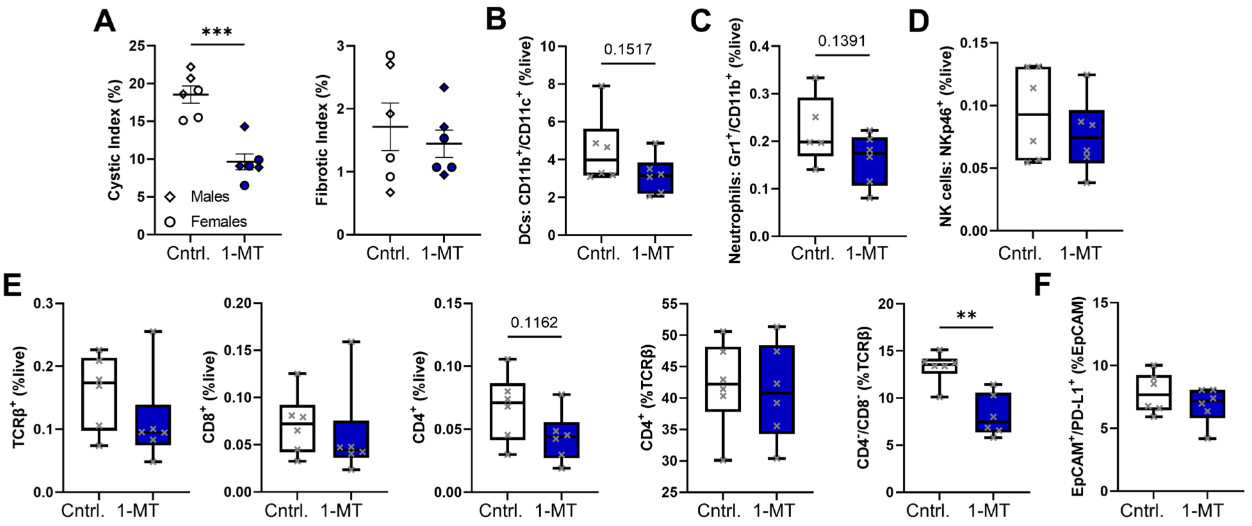


**Supplemental Figure 5 | 1-MT treatment in *Pkd1*$^{RC/RC}$ mice results in less severe cystic disease and associates with changes to the innate and adaptive immune microenvironment.** Quantification of (A) cystic index and (B) fibrotic index in *Pkd1*$^{RC/RC}$ mice treated with (blue) or without (white, Cntrl.) 1-MT. While cystic index was significantly reduced upon treatment, the level of fibrosis was not altered. Quantification of flow cytometry data obtained from kidney single cell suspensions of *Pkd1*$^{RC/RC}$ animals treated with (blue) or without (white, Cntrl.) 1-MT highlights a trend towards reduced numbers of (B) dendritic cells and (C) neutrophils but not (D) natural killer cells. (E) Overall T cell numbers (TCRβ$^{+}$) or their subtypes did not change significantly within *Pkd1*$^{RC/RC}$ kidneys of treated (blue) versus untreated (white, Cntrl.) mice, but the percent of double negative T cells in respect to all T cells did decrease significantly. (F) Expression of the immune checkpoint ligand PD-L1 trends towards a reduction in treated (blue) versus untreated (white, Cntrl.) on *Pkd1*$^{RC/RC}$ EpCAM$^{+}$ cells. N= 3males/3females per treatment group. Statistics: Graphs: (A) mean ± SEM, (B-F) box plot, whiskers 10-90$^{th}$ percentile; Analyses: unpaired t test. P **<0.01, ***<0.001, comparisons with non-significant statistics are not shown.

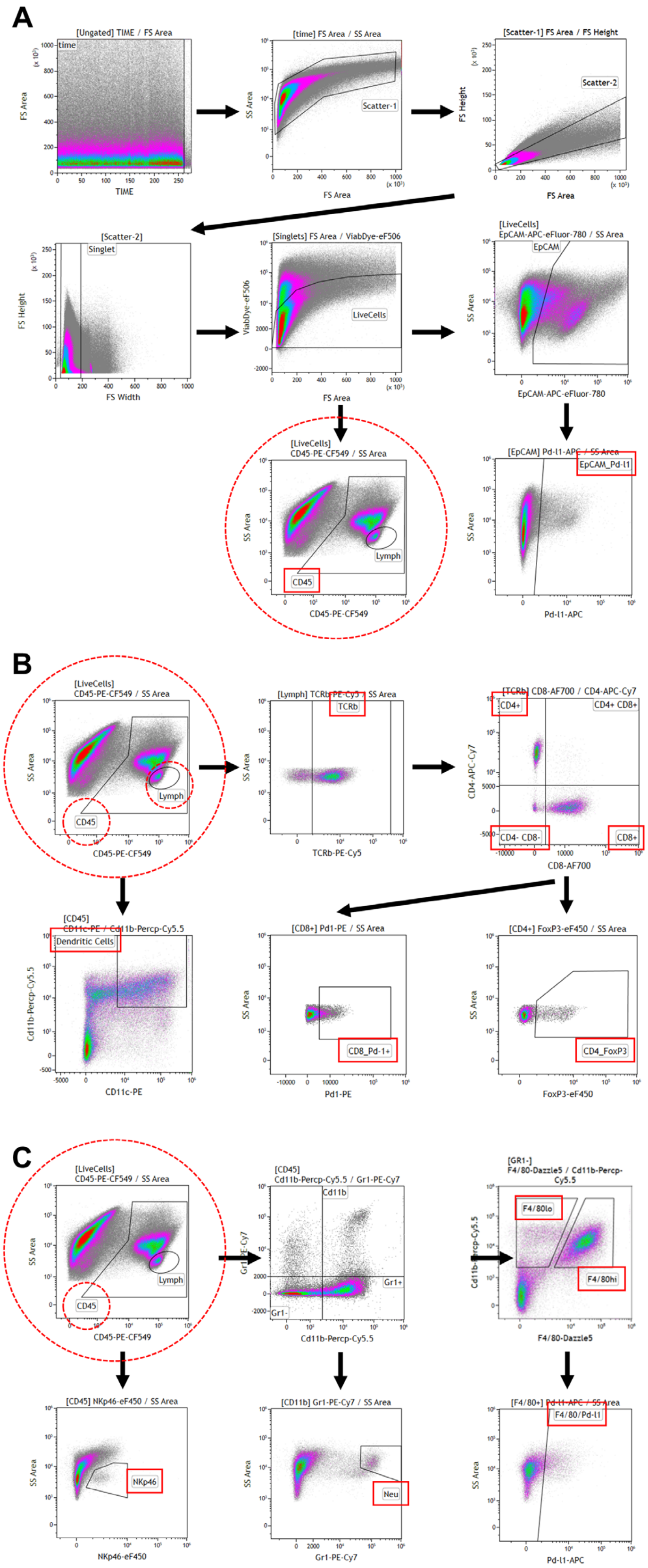


**Supplemental Figure 6 | Gating strategy for flow cytometry analysis of all analyzed cellular subtypes.** Representative flow images are from 3- and 6-month-old C57Bl/6J *Pkd1*$^{RC/RC}$ mice, taken from three different mice. (A) Gating strategy to identify live, CD45$^{+}$, singlets and EpCAM$^{+}$;PD-L1$^{+}$ live singlets. (B) Gating strategy identifying different T cell subtypes and CD8$^{+}$;PD-1$^{+}$ expression, starting with a lymphocyte size exclusion gate, as well as CD45$^{+}$;CD11b$^{+}$;CD11c$^{+}$ dendritic cells starting with the CD45$^{+}$ gate. (C) Gating strategy identifying different innate immune cell populations and their PD-L1$^{+}$ expression starting with the CD45$^{+}$ gate.

### References

1. Research IoLA. *Guide for the Care and Use of Laboratory Animals.* Washington (DC): National Academies Press; 2011.
2. Arroyo J, Escobar-Zarate D, Wells HH, Constans MM, Thao K, Smith JM, et al. The genetic background significantly impacts the severity of kidney cystic disease in the Pkd1(RC/RC) mouse model of autosomal dominant polycystic kidney disease. *Kidney Int.* 2021.
3. Hopp K, Ward CJ, Hommerding CJ, Nasr SH, Tuan HF, Gainullin VG, et al. Functional polycystin-1 dosage governs autosomal dominant polycystic kidney disease severity. *The Journal of clinical investigation.* 2012;122(11):4257-73.
4. Kleczko EK, Marsh KH, Tyler LC, Furgeson SB, Bullock BL, Altmann CJ, et al. CD8(+) T cells modulate autosomal dominant polycystic kidney disease progression. *Kidney international.* 2018;94(6):1127-40.
5. Gainullin VG, Hopp K, Ward CJ, Hommerding CJ, and Harris PC. Polycystin-1 maturation requires polycystin-2 in a dose-dependent manner. *J Clin Invest.* 2015;125(2):607-20.
6. Yuan M, Breitkopf SB, Yang X, and Asara JM. A positive/negative ion-switching, targeted mass spectrometry-based metabolomics platform for bodily fluids, cells, and fresh and fixed tissue. *Nat Protoc.* 2012;7(5):872-81.
7. Baliga MM, Klawitter J, Christians U, Hopp K, Chonchol M, Gitomer BY, et al. Metabolic profiling in children and young adults with autosomal dominant polycystic kidney disease. *Sci Rep.* 2021;11(1):6629.
8. Zhu W, Stevens AP, Dettmer K, Gottfried E, Hoves S, Kreutz M, et al. Quantitative profiling of tryptophan metabolites in serum, urine, and cell culture supernatants by liquid chromatography-tandem mass spectrometry. *Anal Bioanal Chem.* 2011;401(10):3249-61.

9. Chong J, Soufan O, Li C, Caraus I, Li S, Bourque G, et al. MetaboAnalyst 4.0: towards more transparent and integrative metabolomics analysis. *Nucleic Acids Research.* 2018;46(W1):W486-W94.
